# The chemical synthesis of knob domain antibody fragments

**DOI:** 10.1101/2021.06.16.448769

**Authors:** Alex Macpherson, James R. Birtley, Robert J. Broadbridge, Kevin Brady, Yalan Tang, Callum Joyce, Kenneth Saunders, Gregory Bogle, John Horton, Sebastian Kelm, Richard D. Taylor, Richard J. Franklin, Matthew D. Selby, Maisem Laabei, Toska Wonfor, Adam Hold, Douangsone Vadysirisack, Jiye Shi, Jean van den Elsen, Alastair D.G. Lawson

## Abstract

Cysteine-rich knob domains found in the ultralong complementarity determining regions of a subset of bovine antibodies, are capable of functioning autonomously as 3-6 kDa peptides. While they can be expressed recombinantly in cellular systems, in this paper we show that knob domains are also readily amenable to chemical synthesis, with a co-crystal structure of a chemically synthesised knob domain in complex with antigen showing structural equivalence to the biological product. For drug discovery, following immunisation of cattle, knob domain peptides can be synthesised directly from antibody sequence data, combining the power and diversity of the bovine immune repertoire with the ability to rapidly incorporate non-biological modifications. We demonstrate that, through rational design with non-natural amino acids, paratope diversity can be massively expanded, in this case improving the efficacy of an allosteric peptide. As a potential route to further improve stability, we also performed head-to-tail cyclisation, exploiting the unusual proximity of the N- and C-termini to synthesise functional, fully cyclic antibody fragments. Lastly, we highlight the stability of knob domains in plasma and, through pharmacokinetic studies, use palmitoylation as a route to extend the plasma half-life of knob domains *in vivo*. This study presents an antibody-derived medicinal chemistry platform, with protocols for solid-phase synthesis of knob domains; together with characterisation of their molecular structures, *in vitro* pharmacology and pharmacokinetics.

## Introduction

Around 10 % of bovine IgM and IgG antibodies are endowed with ultralong CDRH3^1^, where a knob domain, a series of mini loops stabilised by 2-5 disulphide bonds, is held atop a β-ribbon stalk, over 40 Å from the neighbouring CDRs^2–4^. This striking motif arises as a consequence of limited immune gene segment diversity and is aided by priming mechanisms to introduce cysteine residues during somatic hypermutation^5^.

We have previously shown that knob domains can function autonomously to bind antigen independently of the bovine antibody scaffold^6, 7^. This creates antibody fragments of just 3-6 kDa, so small as to be considered peptides. One notable advantage is that, through immunisation, the vast diversity of the bovine immune system can be exploited to produce high affinity and structurally complex peptides.

Small antibody fragments have previously been isolated from sharks and camelids, the VNAR^8^ and VHH^9^ respectively, which are attractive due to their small paratopes and, in the case of certain VHHs, long CDR loops^10^. Due to their size and structural complexity, recombinant antibody production is typically reliant on DNA engineering and mammalian or microbial cellular machinery. Conversely, A functional, full-length anti-HER2 VHH has been produced by solid-phase synthesis^11^. However, due to the inefficiency of performing solid-phase peptide synthesis on long polypeptide chains, this approach is exceptional.

Chemical synthesis of antibody fragments would be advantageous in certain situations. For example, for antibody fragments where the Fc domain is absent and binding to serum albumin has not been engineered, extension of the plasma half-life (t_1/2_) by the neonatal Fc receptor is impossible^12^. For fragments of less than 60 kDa, renal clearance will markedly reduce exposure^13^. While protease instability can further reduce active concentrations of peptidic or antibody fragment-based drugs.

Medicinal chemists employ a range of peptide design strategies to attenuate renal clearance and extend t_1/2_, such as increasing molecular weight by PEGylation^14^ or conferring reversible binding to serum albumin through conjugation of fatty acids, such as palmitic acid^15^. Head-to-tail cyclisation of peptides is an established approach to limit proteolysis^16^, as is stabilisation of tertiary structure by disulphide bonds^17^; a feature which is already inherent to cysteine-rich knob domains. For drug manufacture, working independently of cell systems may lower manufacturing costs^18^, while the stability conferred by an abundance of disulphide bonds may remove the requirement for a temperature-controlled supply chain.

The first knob domain peptides were isolated through bovine immunisation with human complement C5 protein^6^. C5 is a 188 kDa soluble glycoprotein of the complement cascade and the target of the approved monoclonal antibodies eculizumab^19^ and ravulizumab^20^. It remains the focus of intensive therapeutic research, with further antibodies^21^, peptides^22^, aptamers^23^ and small molecules^24^ in clinical and pre-clinical development^25^, with a view to treating complement induced inflammation and autolysis.

Of the four knob domains which we reported to tightly bind C5, three were functionally modifying and two were demonstrably allosteric (as defined by partial antagonism at asymptotic concentrations)^7^. Notably, one knob domain, K92, demonstrated that selective allosteric inhibition of the alternative pathway can be achieved through C5^7^. Co-crystal structures of the C5-knob domain complexes highlighted the molecular interactions underpinning binding and showed that the knob domain peptides adopted 3-strand β-sheet topologies and were constrained by varying numbers of disulphide bonds^7^. Broadly similar folds are endorsed by the cysteine-rich defensin^26^ and various venom peptides^27^ which are ubiquitous as tools of innate immunity and defence, across all clades, and which are also the focus of drug discovery efforts^28, 29^.

Here, we produce previously reported C5 modifying knob domain peptides^6, 7^ by chemical synthesis and explore their biological activity, structure and *in vivo* pharmacokinetics. This study presents antibody-derived medicinal chemistry as a simple approach to modify knob domains to aid the discovery of drug candidates.

## Results

### Synthesis of bovine knob domains

Four knob domains have previously been reported as tightly binding C5 when expressed recombinantly in a HEK293 cell line and purified from a cleavable protein scaffold: K8, K57, K92 and K149, with each peptide stabilised by two or three disulphide bonds (Figure 1B)^7^. By surface plasmon resonance (SPR), we measured equilibrium dissociation constants (K_D_) for binding of these recombinantly expressed peptides to human C5 (17.8 nM for K8, 1.4 nM for K57, <0.6 nM for K92, and 15.5 nM for K149)^6^.

**Figure 1.**
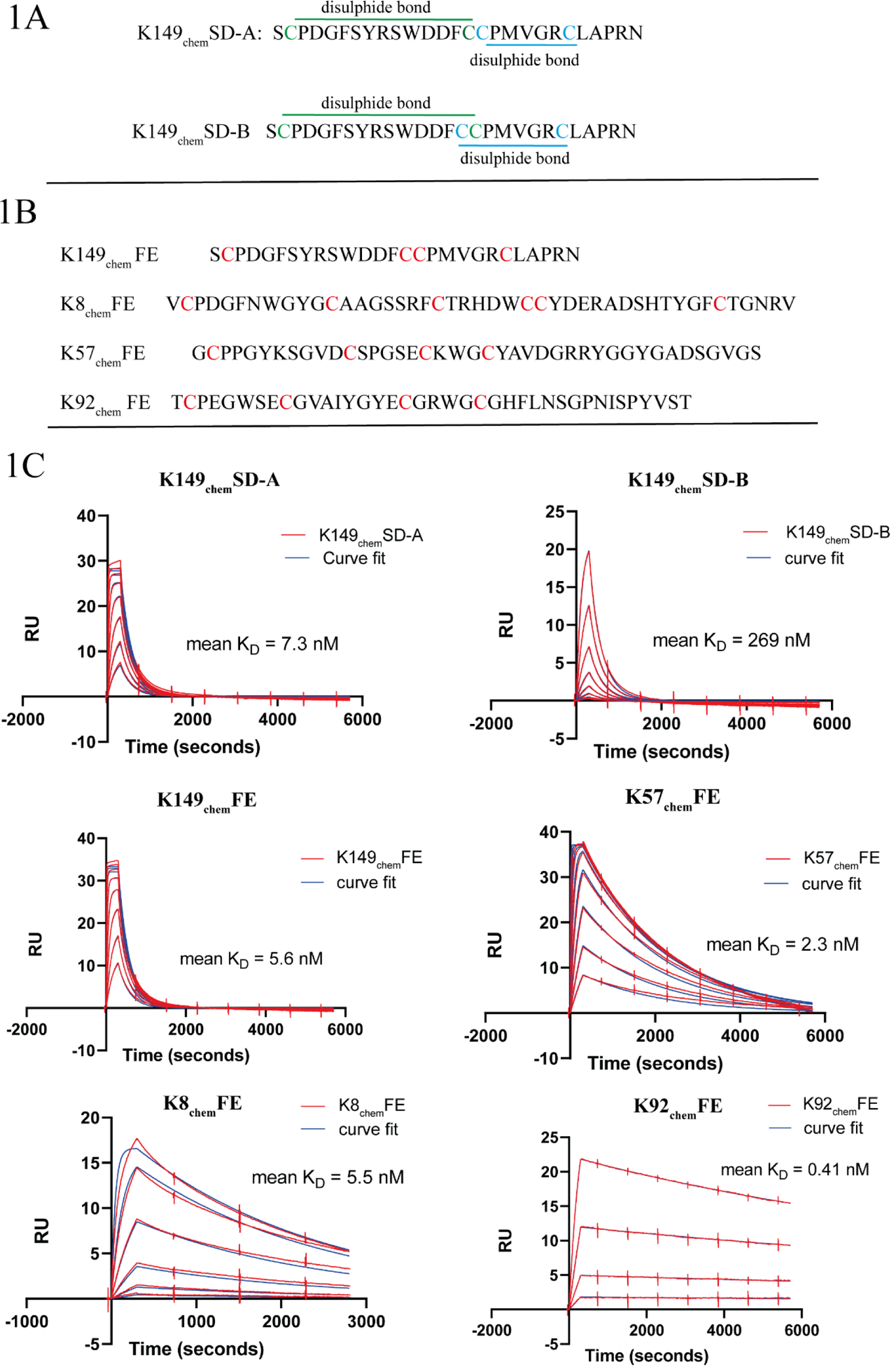
Sequences and binding of chemical knob domain peptides. The sequence and disulphide bond arrangements of K149_chem_SD-A and K149_chem_SD-B, which were synthesised with site specific disulphide bond formation, are shown in panel 1A. The sequences of the peptides produced by refolding under free energy are shown in panel 1B. Panel 1C shows example senrorgrams from a SPR multi-cycle kinetics experiment, with mean equilibrium dissociation constants (K_D_) from *n=3* experiments.

In this study we performed solid-phase peptide synthesis by two methods: firstly, a site-directed method (henceforth suffixed _chem_SD), whereby cysteines in the K149 peptide were specifically protected and deprotected to form disulphide bonds in a preordained manner; and secondly, using a free energy method (henceforth suffixed _chem_FE), where, for all four peptides, thermodynamic-controlled air oxidation was used to obtain the minimum energy form of the disulphide bonds.

For peptides produced by both methods, liquid chromatography/mass spectrometry (LC/MS) confirmed that purities were in excess of 95 % and that masses consistent with the predicted amino acid sequences were unanimously present. A complete list of the knob domain peptide sequences is shown in S1, with accompanying LC/MS characterisation shown in S2.

Binding of knob domain peptides to human C5 was measured by SPR, using a multi-cycle kinetics method. The chemical knob domains produced by the free energy method bound C5 with high affinity (Figure 1C and Table 1), equivalent to values previously reported for the purified recombinant forms of the peptides^6^. For K149, which has two disulphide bonds, the adjacent cysteines C_15_ and C_16_ are unable to pair, giving rise to only two potential disulphide bonding arrangements: K149_chem_SD-A (C_2_-C_15_, C_16_-C_22_) and K149_chem_SD-B (C_2_-C_16_, C_15_-C_22_), shown in Figure 1A. Both forms were synthesised in a site directed manner, and, while both bound C5, K149_chem_SD-B displayed approximately 35-fold lower affinity, with a markedly slower on rate. When produced by the free energy method, K149_chem_FE bound C5 with equal affinity to the higher affinity K149_chem_SD-A form.

**Table 1.** Summary of Biacore multi-cycle kinetics, data from *n=3* experiments.

| **knob domain** | **mean $K_{on}$** | **$K_{on}$ SE** | **mean $K_{off}$** | **$K_{off}$ SD** | **mean $K_D$** | **mean stoich.** |
| --- | --- | --- | --- | --- | --- | --- |
|  | **($M s^{-1}$)** |  | **($s^{-1}$)** |  | **(M)** | **ratio\*** |
| K149<sub>chem</sub>SD-A | 7.13E+05 | 1.08E+05 | 4.84E-03 | 1.08E-03 | 7.31E-09 | 0.82 |
| K149<sub>chem</sub>SD-B | 1.56E+04 | 9.53E+01 | 4.17E-03 | 8.02E-04 | 2.69E-07 | 0.81 |
| K149<sub>chem</sub>FE | 8.61E+05 | 1.60E+05 | 4.48E-03 | 1.57E-03 | 5.57E-09 | 0.85 |
| K57<sub>chem</sub>FE | 2.60E+05 | 2.71E+04 | 5.79E-04 | 1.14E-04 | 2.28E-09 | 0.65 |
| K8<sub>chem</sub>FE | 7.05E+04 | 8.23E+02 | 3.85E-04 | 3.25E-05 | 5.46E-09 | 0.25 |
| K92<sub>chem</sub>FE | 2.83E+05 | 3.79E+04 | 1.01E-04 | 6.32E-05 | 4.11E-10 | 0.40 |
| K8<sub>chem</sub>FE<sup>cyclic</sup> | 9.40E+04 | 1.19E+04 | 2.59E-03 | 8.01E-04 | 2.85E-08 | 0.28 |
| *K57<sub>chem</sub>FE<sup>cyclic</sup> | 1.95E+05 | 2.66E+03 | 3.71E-04 | 1.71E-06 | 1.91E-09 | 0.47 |
| *K92<sub>chem</sub>FE<sup>cyclic</sup> | 1.33E+04 | 2.81E+02 | 2.18E-04 | 1.51E-05 | 1.63E-08 | 0.42 |
| *K149<sub>chem</sub>FE<sup>cyclic</sup> | 3.93E+06 | 1.07E+05 | 4.68E-03 | 9.44E-05 | 1.19E-09 | 0.45 |
| K92<sub>chem</sub>FE A12N | 4.93E+05 | 4.41E+04 | 3.24E-04 | 2.02E-05 | 1.90E-09 | 0.45 |
| K92<sub>chem</sub>FE A12K(Ac) | 1.08E+05 | 1.85E+03 | 3.02E-03 | 2.22E-04 | 5.35E-08 | 0.51 |
| K92<sub>chem</sub>FE A12d-W | 9.24E+03 | 9.60E+02 | 5.85E-04 | 2.97E-05 | 6.81E-08 | 0.46 |
| K57<sub>chem</sub>FE -Palmitoyl | 5.11E+04 | 4.63E+03 | 4.77E-04 | 1.42E-04 | 9.52E-09 | 1.12 |

\*mean stoichiometric ratio is calculated by dividing the measured $R_{max}$ value by the theoretical $R_{max}$

As mispairing of K149 disulphide bonds was tolerated to a certain extent, we next tested the effect of removing disulphide bonds entirely, through reduction and capping of cysteines with iodoacetamide (IAM). Following high resolution LC/MS analysis, to confirm uniform capping had occurred (shown in S3), binding to C5 was again measured by SPR (shown in S4). For K149_chem_FE, reduction and capping of cysteines entirely abrogated binding, while for K92_chem_FE, it resulted in a substantial drop in affinity from 411 pM to 2 µM, predominantly mediated by a decrease in on rate, potentially due to a loss of tertiary structure. Removal of disulphide bonds in K57_chem_FE also affected binding affinity but a reasonable K_D_ of 76 nM was retained.

### Modulation of C5 activation by chemical knob domains

Having demonstrated binding to C5, biological function was evaluated in a range of complement ELISAs and erythrocyte haemolysis assays that were specific for either alternative (AP) or classical pathway (CP) activation. In complement activation ELISAs, which tracked C5b-6, a complex formed from the an activated product of C5, K57_chem_FE was a potent and fully efficacious inhibitor of both the CP and AP (Figure 2E and 2F); K8_chem_FE was a partial inhibitor of the CP and AP (Figure 2A); while K92_chem_FE partially inhibited the AP and showed slight, dose dependent enhancement of the CP (Figure 2C). Consistent with earlier studies with biologically derived K149^7^, K149_chem_FE was a non-functional, silent binder of C5 (data not shown).

**Figure 2.**
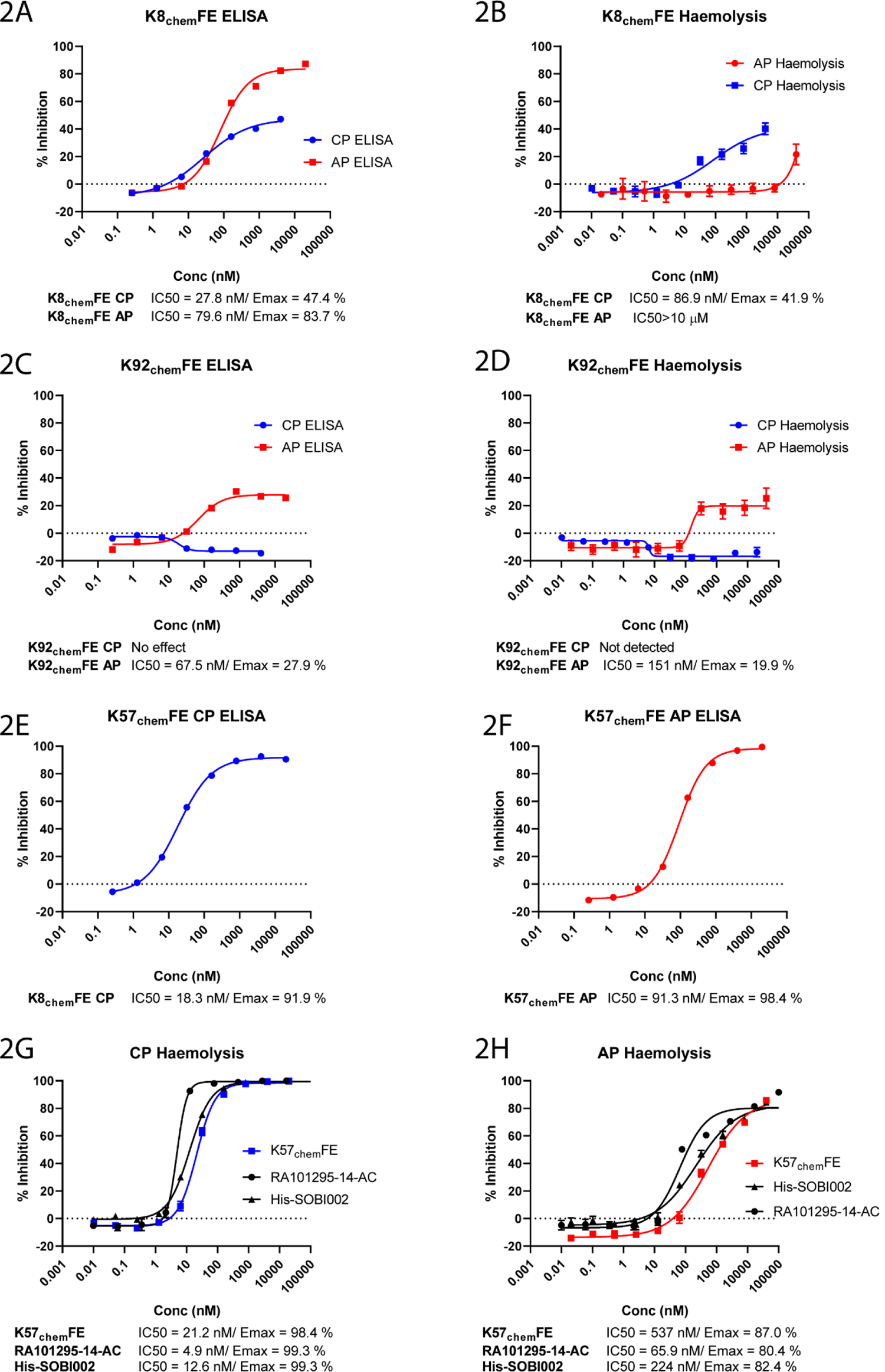
Functional characterisation of chemical knob domains. ELISA and haemolysis data for K8_chem_FE, in both CP and AP driven assays, are shown in panels 2A and 2B, and for K92_chem_FE in panels 2C and 2D. While K8_chem_FE is a partial antagonist, K92_chem_FE is a selective partial antagonist of the AP, consistent with earlier studies using recombinant knob domains. By ELISA, K57_chem_FE is a potent and fully efficacious inhibitor of both pathways, shown in panels 2E and 2F. In panels 2G and 2H, K57_chem_FE is compared to two previously characterised C5 inhibitors, RA101295 and SOBI002, in AP and CP driven erythrocyte haemolysis assays. ELISA and haemolysis data are from *n=3* and *n=6* replicates, respectively.

The peptides were counter screened using ELISAs which measured deposition of C3d, an activation product from complement C3, in response to AP and CP activation (shown in S5). A lack of activity in these assays confirmed that the chemical knob domain peptides did not affect activation of either of the human C5 homologs, C3 or C4, which are both upstream of C5 in the complement cascade.

Behaviour in erythrocyte haemolysis assays was consistent with the ELISAs. K57_chem_FE was a potent and fully efficacious inhibitor of complement mediated cell lysis, K92_chem_FE was active solely in the AP-driven haemolysis assay (Figure 2D) and K8 was a partial inhibitor for the CP and weakly active in the AP assay (Figure 2B). Importantly, these observations in the ELISA and haemolysis assays closely mirror those previously reported with the biological forms of the peptides^7^. In haemolysis assays, K57_chem_FE was broadly equivalent to two previously characterised C5 inhibitors (Figure 2G and 2H): RA101295^30^, a close analogue of the UCB-Ra Pharma macrocyclic peptide Zilucoplan^31^, which is currently in phase III trials, and SOBI002^32, 33^, an affibody from Swedish Orphan Biovitrum, which was discontinued after showing transient adverse effects in a phase 1 trial^34^. While the three inhibitors display equivalent efficacy, RA101295 exhibited approximately ten-fold higher potency than K57_chem_FE and SOBI002.

### Elucidation of the molecular structure of K92**_chem_**FE in complex with C5

To permit comparison of the molecular structure of a chemically synthesised knob domain, a crystal structure of K92_chem_FE in complex with C5 was determined at a resolution of 2.57 Å (Figure 3A, data collection and refinement statistics are shown in Table 2). The C5-K92_chem_FE complex was crystallised under the same conditions previously reported for the C5-biological K92 (K92_bio_) complex (PDB accession: 7AD6)^7^. A stringent mFo-DFc simulated annealing omit map of the C5-K92_chem_FE complex, with the peptide deleted from the model, shows clear electron density for the peptide at 1.0 sigma (Figure 3C), while the final structure shows the fold and disulphide bond arrangement of K92_chem_FE to be contiguous to K92_bio_ (Figure 3B). Analysis with the macromolecular structure analysis tool PDBPiSA^35^, confirmed that the molecular interactions which sustain binding to C5 are consistent between K92_chem_FE and K92_bio_ (Shown in S6).

**Figure 3.**
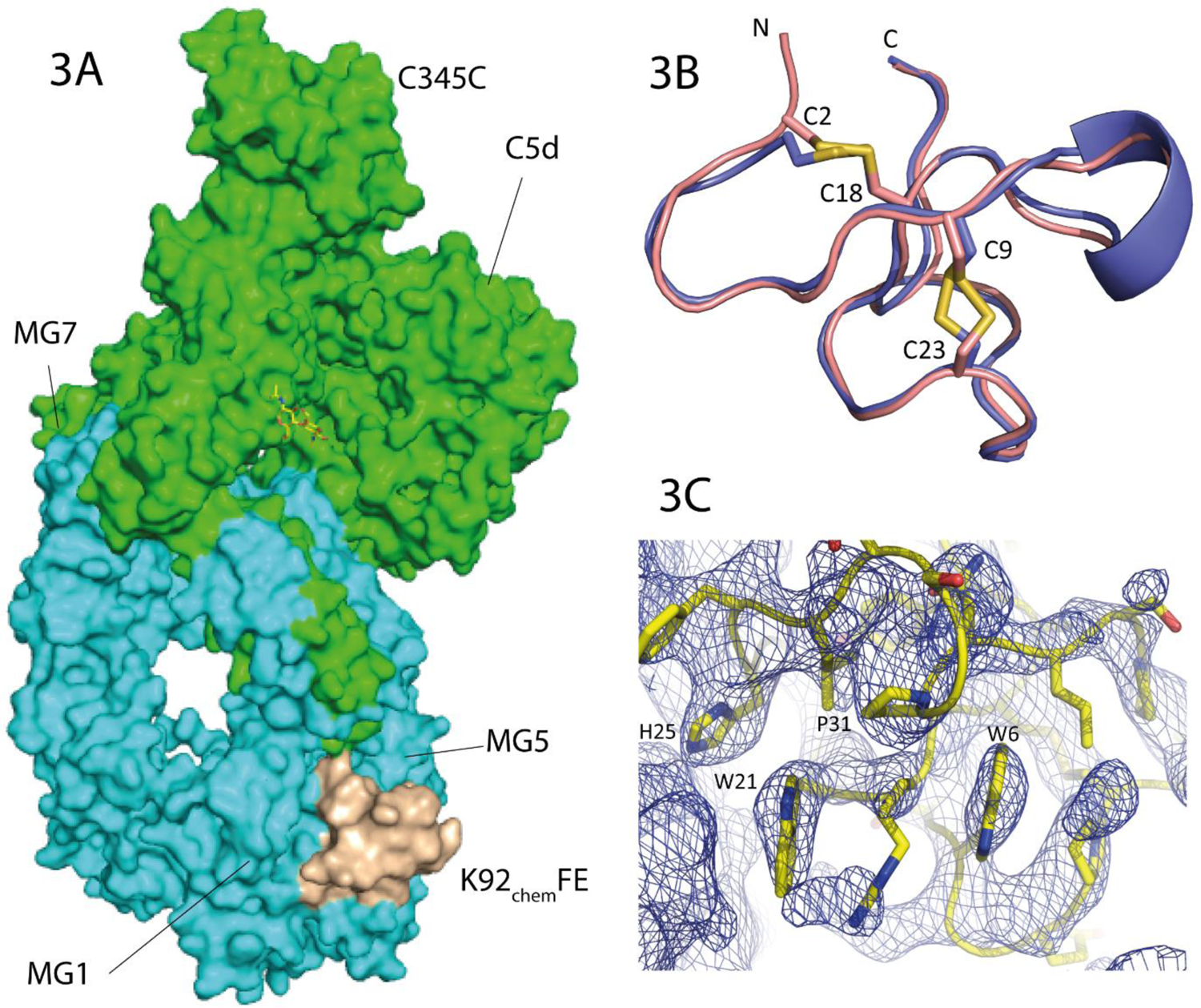
Crystal structure of the C5-K92_chem_ complex. Panel 3A shows K92_chem_FE in complex with C5, binding to an epitope located between the MG1 and MG5 domains. Panel 3B shows an overlay of the recombinant and chemically synthesised K92 peptides. K92_chem_ (shown in wheat) and the K92 recombinant peptide (cyan) were superposed (RMSD = 0.55Å) and shown as a backbone trace. N- and C-termini are highlighted as are the disulphide bonds arrangements with the cystines shown as sticks. Panel 3C shows a simulated annealing omit map of the K92_chem_FE peptide. The blue mesh shows a mFo-DFc simulated annealing omit map calculated in PHENIX and contoured at 1.0 sigma around K92_chem_FE. In the omit calculation the peptide was deleted from the model. The peptide is displayed in yellow in ribbon representation with side chains shown as sticks and coloured according to the atom type (nitrogen in blue, oxygen in red and carbon in yellow). Selected side chains are highlighted.

**Table 2.**
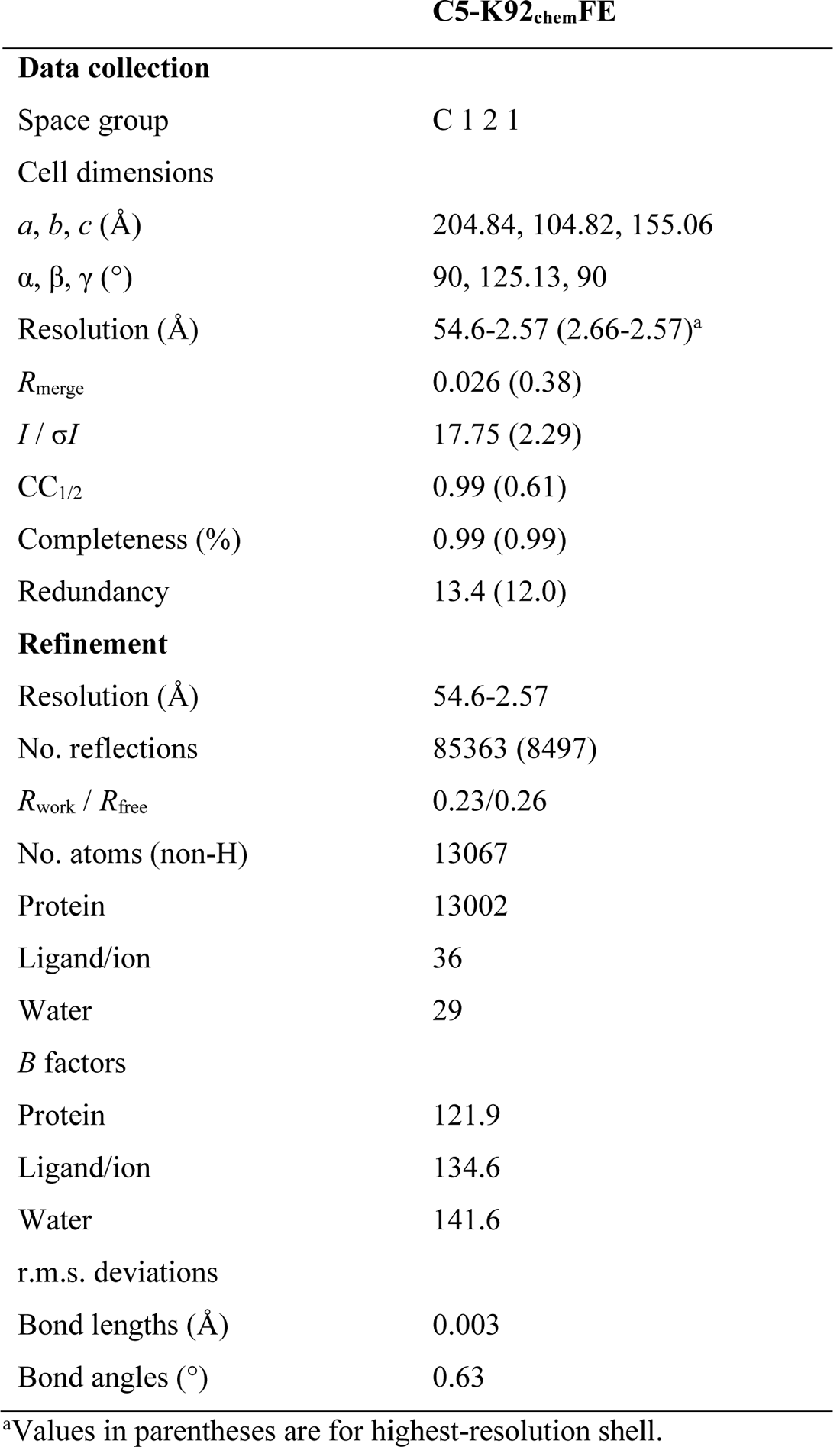
Data collection and refinement statistics (molecular replacement).

### Incorporation of non-natural amino acids enables structure-based antibody drug design, akin to small molecule medicinal chemistry

Examination of the K92_chem_FE binding interface with C5 revealed that electrostatic interactions were comparatively sparse, with the structure suggesting just ten hydrogen bonds were present, which were predominantly mediated via the polypeptide backbone. We noted a bifurcated pocket on the β-chain of C5, adjacent to the residues A12 and I13 of K92_chem_FE, and attempted to rationally design mutations to enable new interactions to be made with C5 within this region, with the goal of improving the biology efficacy. Residue A12 offered scope to contact the α-chain on C5 via N805_C5_ (numbering based on the mature C5 sequence), which the crystal structure of the complex suggested did not have strong non-covalent molecular interactions with K92, as well as access a cavity on the C5 β-chain, which is flanked by residues L152_C5_, Q532_C5_ and V534_C5_ (Figure 4A). We therefore designed mutants for A12, using Molecular Operating Environment’s (MOE, version 2019^36^) residue scan function (shown in S7 and S8). We predominantly focused on non-natural and D-forms of amino acids, which have been shown to be beneficial in improving the stability, potency, permeability, and bioavailability of peptide-based therapies^37^.

**Figure 4.**
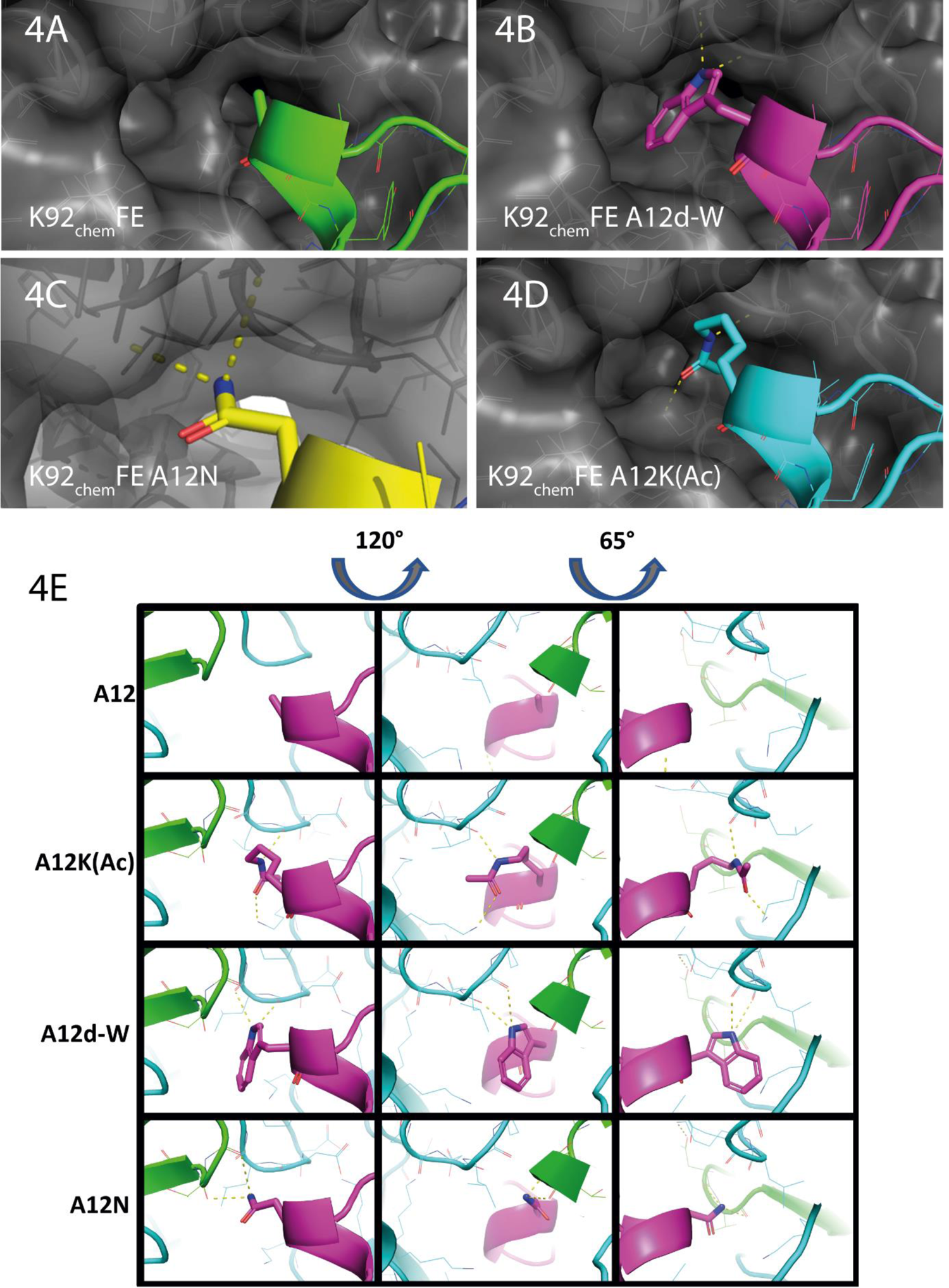
Rational design with non-natural amino acids. Panel 4A shows the location of alanine 12 and an adjacent cavity, while panels 4B, 4C and 4D show predicted binding poses for mutants comprising the natural amino acid asparagine (A12N) as well as the D-form of tryptophan (A12d-W) and an acetylated lysine [A12K(Ac)]. Panel 4E shows the predicted binding poses and interactions from three different angles.

The final eight designs, shown in S9, contained mono and bicyclic side chains, such as D-tryptophan, benzothiazole and 2-oxo-histidine; acetylated and methylated lysine; L- and D-forms of ornithine, as well as the natural amino acid asparagine. The mutants were tested in SPR multi-cycle kinetics experiments and in AP and CP complement ELISAs. While all the mutants were readily synthesised, the assays identified three mutations of note: asparagine (A12N [shown in Figure 4C and 4E]), D-tryptophan (A12d-W [shown in Figure 4B and 4E]) and acetylated lysine (A12K[Ac]), shown in Figure 4D and 4E.

Our SPR data (Table 1 and Figure 5E-H), show that we did not improve binding affinity relative to K92_chem_FE in any of these mutants. While the A12N mutation did not result in a loss of affinity, both A12d-W and A12K(Ac) suffered 130 and 166-fold reductions in affinity for C5, with K_D_ values of 53.5 nM and 68.1 nM, respectively. However, while all three mutations resulted in an acceleration in k_off_, for A12N, there was a compensatory improvement in k_on_ (Table 1).

**Figure 5.**
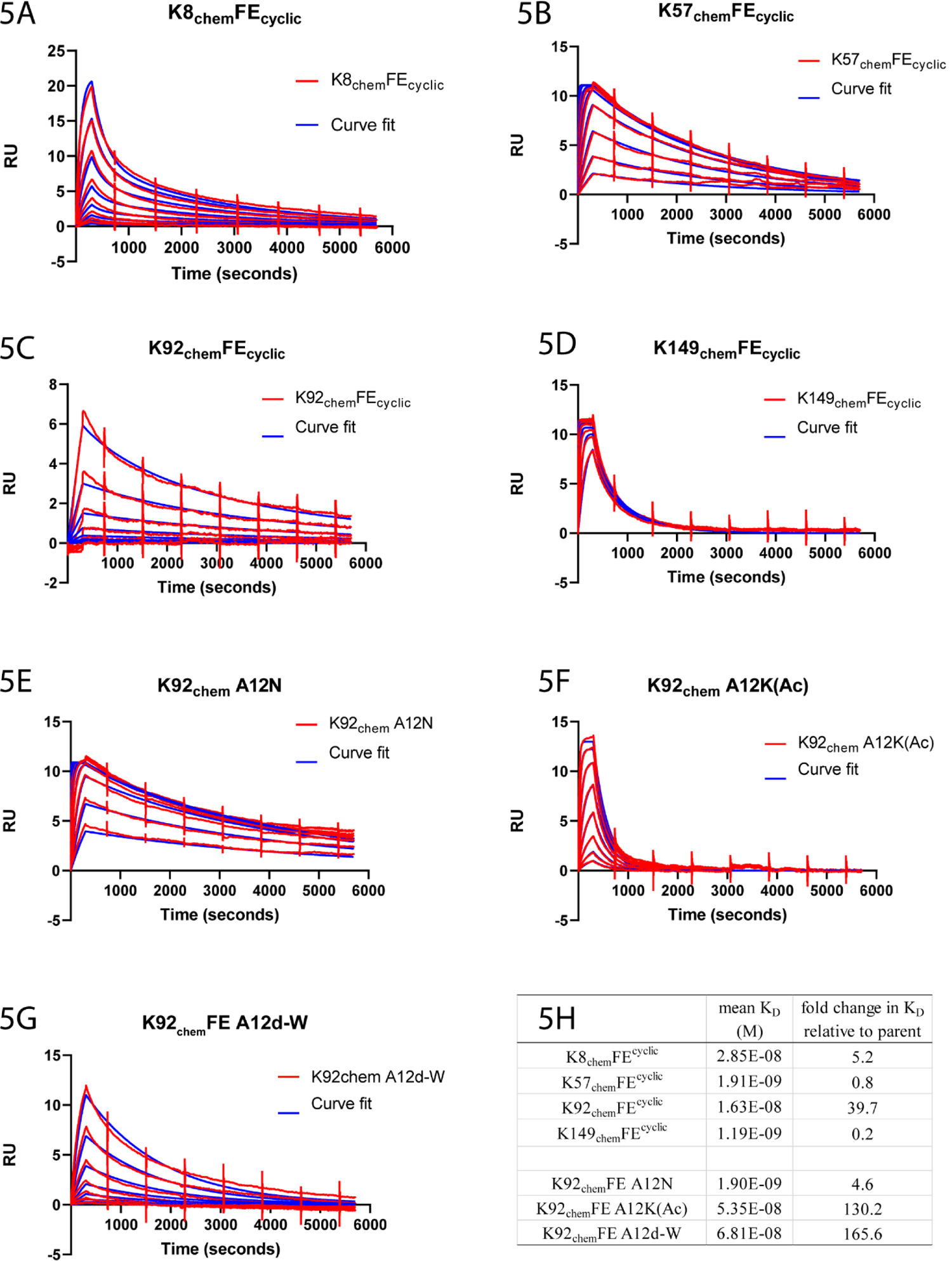
Binding of fully cyclic knob domains and K92_chem_FE mutants. Example sensorgrams from a SPR multi-cycle kinetics experiment, with mean equilibrium dissociation constants (K_D_) from *n=3* experiments, are shown. Fully cyclic knob domains are shown in panels 5A-D and K92_chem_FE alanine 12 mutants are shown in panels 5E-G. The fold change in affinity, relative to the unmodified parent knob domain, is shown in panel 5H.

As shown earlier, K92_chem_FE has no effect on the CP but is demonstrably allosteric in the AP ELISA, with an IC_50_ of 84 nM and an E_max_ of just 27 %. Consistent with the K92_chem_FE parent compound, all mutations had no effect in the CP ELISA (Figure 6F). In the AP ELSA, the A12N mutation gained approximately a log of potency (IC_50_ of 4 nM) but did not affect E_max_ to any great extent (Figure 6G). Similarly, the A12d-W mutant appeared not to have lost potency or efficacy, despite having a markedly lower affinity than the parent (Figure 6I). For drug binding studies *in vitro*, potent compounds frequently have faster k_on_, which speeds their onset of action^38^, subsequently an increase in k_on_ can compensate for a reduction in binding affinity in a short assay, which may explain the gain in potency for the A12N mutant. Conversely, to prolong drug action *in vivo,* the endurance of a drug– target complex is usually more contingent on k_off_.

**Figure 6.**
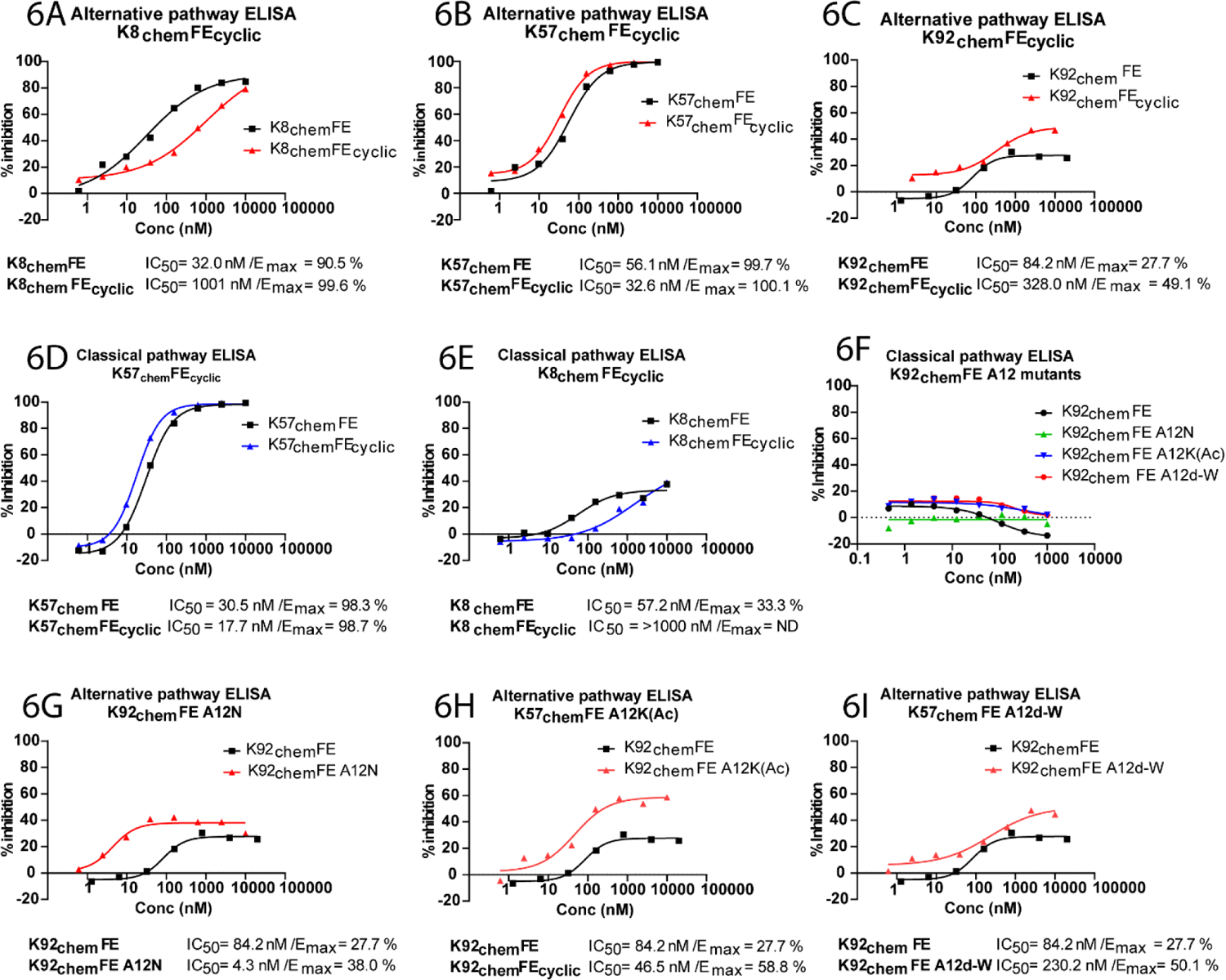
Functional data for fully cyclic knob domains and K92_chem_FE mutants. AP ELISA data for cyclic knob domain peptides are displayed in panels 6A-C and for K92_chem_FE mutants in panels 6G-I. CP ELISA data for K57_chem_FE_cyclic_ and K8_chem_FE_cyclic_ are shown in 6D and 6E, respectively. Panel 6F shows that, similar to the parent compound, K92_chem_FE mutants are inactive in a CP driven ELISA. ELISA data are from *n=3* replicates.

Interestingly, the A12K(Ac) mutant also displayed a slight increase in potency but exhibited a significant increase in E_max_, with a two-fold increase in efficacy observed (Figure 6H). For allosteric compounds, biological efficacy is due to an effect on the protein target’s conformation that is uncorrelated with binding affinity^40^, but a conformationally constrained, or rigidified, compound may be beneficial, particularly for small molecules^41^. While we were unsuccessful in increasing the affinity of the complex, we show that the rational incorporation of non-proteogenic amino acids into knob domains can be a route to modulate biological efficacy and our data suggest that D-amino acids may also be tolerated, which may be beneficial when exploring routes to attenuate proteolysis.

### Synthesis of fully cyclic antibody fragments

For development of peptide drug candidates, head-to-tail cyclisation can extend exposure *in vivo* by preventing exopeptidase cleavage^42^. To create small, cyclic antibody fragments, we synthesised head-to-tail cyclised forms of our knob domains, which are subsequently termed: K8_chem_FE^cyclic^, K57_chem_FE^cyclic^, K92_chem_FE^cyclic^ and K149_chem_FE^cyclic^.

All cyclic knob domains bound C5 (Table 1 and Figure 5A-D). For K57_chem_FE^cyclic^ and K149_chem_FE^cyclic^ affinity was improved, but binding was attenuated in K8_chem_FE^cyclic^ and K92_chem_FE^cyclic^ (Figure 5H). Biological activity was again tested in AP and CP ELISAs. The activity of K57_chem_FE^cyclic^ and K92_chem_FE^cyclic^ was not obviously affected by cyclisation, while K8_chem_FE^cyclic^ displayed a modest loss of potency relative to K8_chem_FE (Figure 6A-E). Consistent with the parent compound, K149_chem_FE^cyclic^ had no discernible effect on complement activation, despite binding C5 (data not shown). Overall, these data suggest that knob domains are amenable to cyclisation and, while some optimisation may be required in certain cases, this approach can be explored to improve stability.

### Pharmacokinetics and *in vitro* plasma stability of bovine knob domain peptides

Finally, we sought to explore the pharmacokinetic profile of chemically synthesised knob domain peptides in rodents. As target binding may influence pharmacokinetics, we first tested for binding to C5 protein from *Rattus norvegicus* (rat). By SPR, K8_chem_FE was cross reactive with rat C5 protein (as measured by single-cycle kinetics, shown in S10), as well as C5 from other species (data not shown), while K57_chem_FE was specific for human C5. To explore the effect of fatty acid conjugation, we synthesised a palmitoylated form of K57_chem_FE, with attachment via a Gly-Ser-Ser-Gly linker at the N-terminus, and determined an apparent four-fold reduction in binding for affinity for human C5 by SPR (Table 1 and S11).

To test if the knob domains were resistant to proteolysis by virtue of their abundant disulphide bonds, we employed HPLC mass spectrometry to track the stability of K8_chem_FE, K57_chem_FE and K57_chem_FE-palmitoyl in human, rat and *Mus musculus* (mouse) plasma at 37 °C, over a period of 24 hours. The unmodified knob domains were predominantly stable in human plasma, with K8_chem_FE not degraded in plasma from humans, mice or rats (Figure 7C). For K57_chem_FE, in excess of 75 % remained intact after 24 hours in human plasma, although markedly more proteolysis was observed in rodent plasma (Figure 7A). The addition of a fatty acid to K57_chem_FE-palmitoyl conferred protection from proteolysis in rodent plasma (Figure 7B), by virtue of binding to serum albumin.

**Figure 7.**
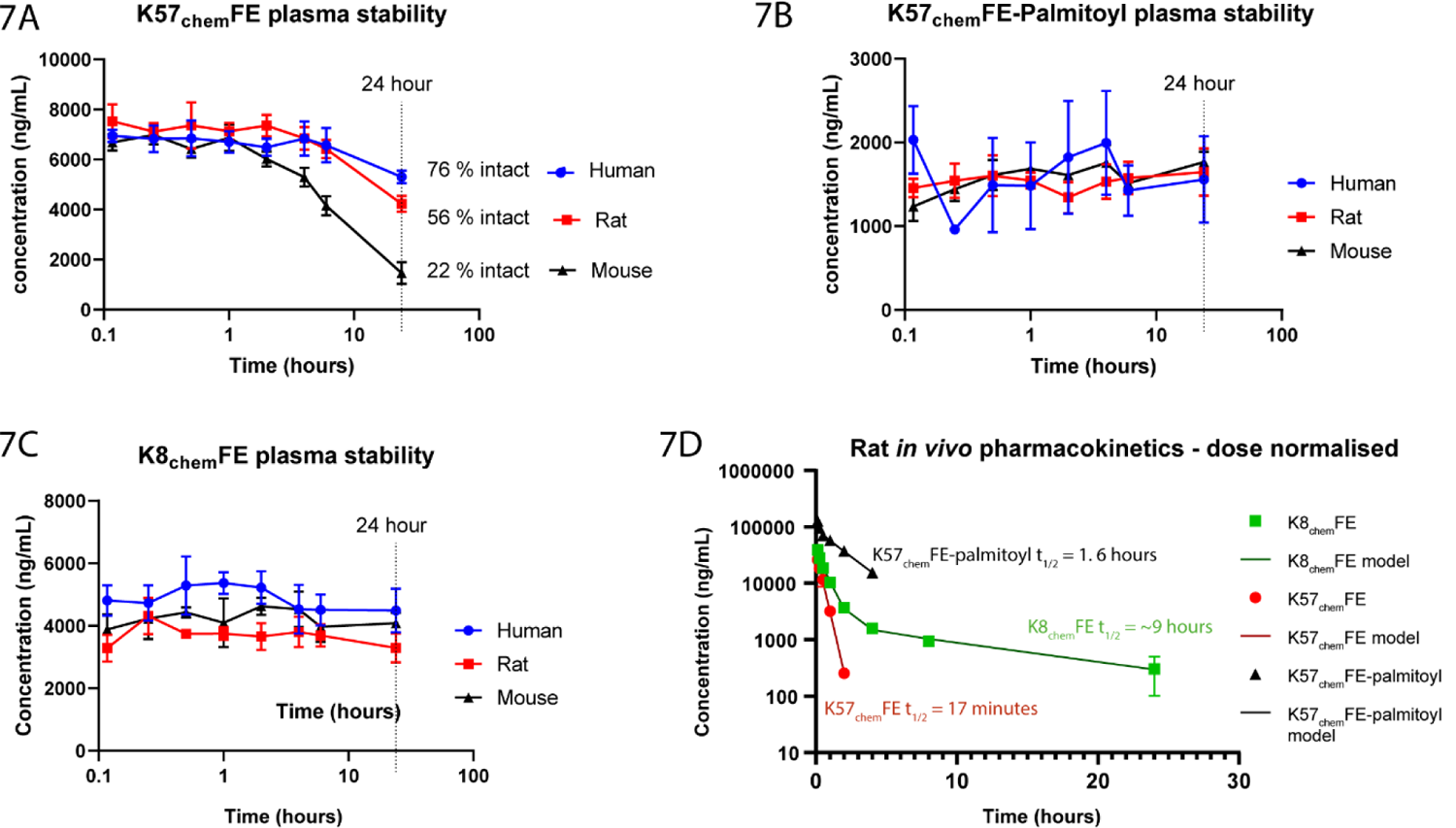
Plasma stability and *in vivo* pharmacokinetics. The plasma stability of K57_chem_FE, K57_chem_FE-Palmitoyl and K8_chem_FE are shown in panels 7A-C. The peptides are stable in plasma for a minimum of ten hours. The pharmacokinetics (PK) of the knob domains are shown in panel 7D, following intravenous dosing to Sprague Dawley rats. Unmodified K57_chem_FE is rapidly cleared but chemical modification with palmitoyl extends exposure. Unmodified K8_chem_FE exhibits a plasma half-life of ∼9 hours, mediated by tight binding to rat C5.

The pharmacokinetics of K8_chem_FE, K57_chem_FE and K57_chem_FE-palmitoyl were measured following dosing via an intravenous bolus to Sprague Dawley rats (Figure 7D). Following administration at 10 mg/kg, K57_chem_FE was eliminated in a rapid manner typical of low molecular weight peptides and proteins (t_1/2_ = 17 mins/ plasma clearance [Clp] = 10.6 ml/min/kg), which is typically due to glomerular filtration in the kidney. In contrast, K8_chem_FE tightly bound rat C5 protein and adopted target-like pharmacokinetics, resulting in markedly improved exposure (t_1/2_ = ∼9 hours/ Clp = 3.3 ml/min/kg). Due to poor solubility, K57_chem_FE-palmitoyl was tested at a lower dose of 1 mg/kg; as expected, the conjugation of a palmitic fatty acid extended exposure, relative to unmodified K57_chem_FE (t_1/2_ = 1.6 hours/ Clp = 0.8 ml/min/kg). This suggests that conjugation of fatty acids, potentially in combination with cyclisation or other chemical modifications, is a viable route to extend the biological exposure of chemical knob domains *in vivo*.

Our data suggest that, despite their biological origin, knob domain peptides are viable chemical entities and that this greatly aids their potential developability as drug candidates. In chemical form they present with high potency and are inherently stable in plasma. Their pharmacokinetics are akin to conventional peptides - moreover, unusually for antibody derived molecules, chemical modification offers a tractable route to improve their pharmaceutical properties.

## Discussion

We have shown, using various examples, formats and modifications, that knob domains are amenable to chemical synthesis. Crucially, the chemically derived peptides bind antigen with high affinity and are active in biological assays, with K57_chem_FE offering a comparative *in vitro* profile to SOBI002 and a peptide with a similar core sequence to zilucoplan, both of which are clinical stage C5 inhibitors.

We present chemical knob domains as the first examples of antibody fragments which are readily amenable to medicinal chemistry approaches. For drug discovery, medicinal chemistry typically focuses on the optimisation of drug potency, pharmacokinetics and biodistribution. Here we exemplified chemical strategies that may be applied to improve the stability and biology efficacy of knob domain peptides.

To modulate biological efficacy, this study used structure-based drug design with non-natural amino acids to access latent interactions which were not employed in the natural paratope. Non-natural amino acids are commonly used to increase the binding affinity of peptides. Compstatin, a macrocyclic inhibitor of complement C3 which, in a modified form, was recently approved for paroxysmal nocturnal haemoglobinuria^43^, is one such example; where incorporation of a hydrophobic 1-methyltryptophan results in a 264-fold increase in activity^44^. Similarly, non-proteinogenic residues have been used in the generation of high affinity human leukocyte antigen blockers, which competitively displace the antigenic peptide on major histocompatibility complex receptors to prevent T-cell recognition of the complex^45^. To improve stability, adjacent D-amino acids are more resistant to degradation by natural proteases than their L-enantiomeric counterparts, and this represents a strategy to employ against proteolysis^37^, through stabilization of backbone conformation and elimination of the cleavage site^46^. For knob domain peptides, this approach is complementary to the optimisation of the natural amino acid sequence, which occurs *in vivo* within the germinal centres of the secondary lymphoid organs when a cow is exposed to an antigen. The rational application of non-natural amino acids, in this post-immune setting, can bring chemical space and physiochemical properties, that are not at the disposal of the bovine immune system, into play.

There are several methods for incorporation of non-natural amino acids into antibody fragments or other small proteins, as part of attempts to either improve affinity, conjugate toxins for antibody drug conjugates^47^ or to introduce chemical handles for photocoupling^48^. These primarily focus on either genetic codon expansion technology to permit incorporation of unnatural amino acids into the translational machinery of the host cell^49^, post translation bis-alkylation of cysteines to the non-natural amino acid Dha^50^, or complete solid-phase synthesis of proteins^11, 51^. While these solutions are elegant, they are restricted; genetic codon expansion can currently only change one residue at a time within a single-engineered expression system, while bis-alkylation of cysteines can only be used to introduce Dha. Chemical synthesis of proteins seems the most attractive option and has been used to produce a functional VHH fragment which bound HER2^11^. With established synthesis methods, this approach is suboptimal due to the inefficiency associated with solid-phase synthesis of long polypeptide chains.

However, new methods in flow chemistry may help to overcome this limitation, with an automated instrument which can manufacture polypeptide chains of up to 164 amino acids, over 327 consecutive reactions, recently reported ^51^.

Our data suggest that the knob domains are highly resistant to plasma proteolysis *in vitro*, potentially due to their network of disulphide bonds. We have shown that the proximity of the N- and C-termini can give rise to fully cyclic antibody fragments, which may provide a route to reduce proteolysis even further, especially when used in conjunction with other chemical approaches.

When dosed *in vivo*, knob domains exhibit typically peptide-like pharmacokinetics and in unmodified form are likely subject to renal clearance, unless target-mediated drug disposition is achieved through tight binding to a target. We have shown that by using chemical synthesis, simple modifications such as palmitoylation can be readily incorporated to attenuate renal clearance and produce lead-like molecules, which are consistent with once daily dosing.

This chemical method may also provide a route to decouple small antibody fragments from the cost constraints associated with manufacturing in cell systems. The cost of antibody therapies has come under scrutiny in the face of global COVID-19 pandemic and alternatives are being sought to address the so called ‘access gap’, whereby 80 % of the global licenced antibody sales are within the United States, Europe and Canada^52^. We hope that the abundance of disulphide bonds and apparent stability of the knob domains may remove the requirement for a cold supply chain, in a similar manner to other peptide drugs.

We present a chemical biology approach for bovine antibody-derived knob domain peptides, demonstrating structural and functional equivalence from biologically expressed and chemically synthesised material. Combining the sequence diversity of immune systems with structure-based medicinal chemistry techniques provides new opportunities to create innovative therapeutic solutions for patients.

## Methods

### Purification of C5 protein

C5 protein was affinity purified using an E141A, H164A OmCI column^53^. Briefly, human and rat sera (TCS biosciences) were diluted 1:1 (v/v) with PBS, 20 mM EDTA, pH 7.4 and applied to a 5 mL Hi-Trap NHS column (GE Healthcare), which contained 20 mg of immobilised E141A H164A OmCI protein, at a flow rate of 1 mL/minute. The column was washed with 5x column volumes of PBS, and C5 protein was eluted using 2 M MgCl_2_ and extensively dialysed into PBS.

### Peptide synthesis of knob domains

All peptides were synthesised using solid phase synthesis, employing the Fluorenylmethyloxycarbonyl (Fmoc) technique^54^. Synthesis was performed in a sequential manner in the C to N direction on robotic synthesisers (Symphony, Protein Technologies). The syntheses were initiated upon appropriate polystyrene supports (Nova Biochem) with the first amino acid attached via a Wang linkage to the carboxyl, substitution of 0.3 mM/g. Chain elongation was facilitated by using a twenty minute double coupling strategy with a 3 fold molar excess of reagents to the loading of the resins with N-alpha protected amino acids dissolved in dimethylformamide (DMF) (side chains were orthogonally protected with suitable protecting groups for Fmoc chemistry) and the coupling reagent 2-(1H-Benzotriazole-1-yl)-1,1,3,3-tetramethylaminium tetrafluoroborate (TBTU), in the presence of N,N-diisopropylethylamine (DIPEA). The temporary amino protection was removed by two, 5-minute treatments with 20 % piperidine in DMF. After the peptide sequences were complete the peptidyl resins were treated with a mixture of trifluoroacetic acid (TFA), ethane dithiol and tri isopropyl silane, 95:3:2 for 3 hours to cleave the peptides and all protecting groups. The peptides were isolated by filtration and trituration with diethyl ether. The peptides were dissolved in acetonitrile water and freeze dried before purification.

### Purification and cyclisation of knob domains

It was found that the most expedient and high yielding way to obtain these cyclic peptides with greater than two disulphide bonds was to reversed phase high performance liquid chromatography (RP-HPLC) purify the linear sequence and immediately initiate cyclisation without a freeze-drying step, otherwise much insoluble polymeric material resulted.

Cyclisation was achieved by using thermodynamic-controlled air oxidation to obtain the minimum energy form of the disulphides in the sequence, employing a mixture of reduced and oxidised glutathione.

Crude peptides were dissolved in dimethyl sulfoxide (DMSO)/ water and treated with tris (2-carboxyethyl) phosphine (TCEP) to ensure the disulphide bonds were fully reduced. The peptides were purified by RP-HPLC, using a Varian Prostar system equipped with two 210 pumps and a 355 UV spectrophotometer. Running buffers were for pump A, solvent A, 0.1% (v/v ammonium acetate in water, pH 7.5 −7.8), and for pump B, solvent B, 100% acetonitrile. The peptide was introduced to a prep RP-HPLC column (C18 Axia, 22 mm x 250 mm, 5 micron particle, size 110 angstrom pore size, Phenomenex). The linear sequence was eluted from the column by running a gradient between solvents A and B, 5% B to 65% B over 60 minutes. Linear peptide was identified by electrospray ionisation mass spectrometry.

The solution of the linear peptide (approximately 50 mL) was added to 500 mL of a cyclisation buffer, (0.2 M phosphate buffer, pH 7.5, containing 1mM EDTA, 5mM reduced glutathione, and 0.5 mM oxidised glutathione). The solution was stirred at room temperature for 48 hours. After which, a small sample was analysed by analytical HPLC to assess the level of cyclisation.

When the cyclisation was deemed sufficiently complete, the whole buffer containing the peptide was pumped onto a preparative RP HPLC column (C-18 Axia as above). The cyclic peptide was eluted using a gradient between solvent A (0.1 % TFA in water) and solvent B (0.1% TFA in acetonitrile) of 5% B to 65% B in 60 minutes. Fractions identified as the correct compound were freeze dried before analysis.

### Synthesis of head-to-tail cyclised knob domains

For head-to-tail cyclisation, we used Gly-Cys as the point of cyclisation and used a thioester strategy employing native chemical ligation. Generation of a thioester upon the c-terminal Gly residue does not result in racemisation; while cyclisation using the purified side chain deprotected thioester improves solubility, giving cleaner products and better yields.

N-terminal tert-butyloxycarbonyl (boc) protected peptides were made by Fmoc peptide synthesis using glycine loaded 2-chlorotrityl resin, with cleaving from the resin via 1% TFA treatment. The peptide benzyl-thioesters were generated through reaction with benzyl-mercaptan and 1-benzotriazole-tris-dimethyl. The resulting protected peptidyl-thioester species underwent a complete side chain cleavage, using the normal TFA scavenger cleavage regime. Finally, the peptides were HPLC purified, and then cyclised head-to-tail via native chemical ligation, (by dissolving the peptides in phosphate buffer, pH 7.8, with a 3-fold molar excess of TCEP present, overnight). The head-to-tail cyclised peptides were freeze dried, and intra-chain disulphide bond formation was again achieved using thermodynamic-controlled air oxidation with a mixture of reduced and oxidised glutathione, to obtain the minimum energy form of the disulphides in the sequence, as described previously.

### SPR multicycle kinetics

Kinetics were measured using a Biacore 8K (GE Healthcare) with a CM5 chip, which was prepared as follows: 1-ethyl-3-(3-dimethylaminopropyl)-carbodiimide (EDC)/ N-hydroxy succinimide (NHS) was mixed at 1:1 ratio (flow rate, 10 µL/min; contact time, 30 seconds), human and rat C5 proteins at 1 ug/mL in pH 4.5 sodium-acetate buffer, were injected over flow cell one only (flow rate, 10 µL/min; contact time, 60 seconds). Final immobilization levels in the range of 2000-3000 response units (RU) were obtained, to yield theoretical Rmax values of ∼50-60 RU. Serial dilutions of peptide were prepared in HBS-EP buffer and injected (flow rate, 30 µL/min; contact time, 240 seconds; dissociation time, 6000 seconds). After each injection, the surface was regenerated with two sequential injections of 2M MgCl_2_ (flow rate, 30 µL/min; contact time, 30 seconds). Binding to the reference surface was subtracted, and the data were fitted to a single site binding model, using Biacore evaluation software.

### Complement ELISA

Assays were run using the CP and AP Complement functional ELISA kits (SVAR, COMPL 300 RUO), as per the manufacturer’s protocol. For sample preparation, serum was diluted as per the respective protocol for the CP and AP assays; serial dilutions of peptides were prepared and allowed to incubate with serum for 15 minutes at room temperature, prior to plating.

### Haemolysis assays

For AP, 50 µL of 24% normal human sera (Complement Technology), 50 µL of 20 mM MgEGTA (Complement Technology), and 48 µL of GVB0 buffer (0.1 % gelatin, 5 mM Veronal, 145 mM NaCl, 0.025 % NaN_3_, pH 7.3, Complement Technology) were aliquoted into a single well of a 96-well tissue culture plate (USA Scientific) then mixed with 2 µL of inhibitors serially diluted in DMSO. Following equilibration for 15 minutes at room temperature, 50 µL of rabbit erythrocytes (Complement Technology), at 2.5 x10^7^ per well, were added to the plates, which were then incubated at 37°C for 30 minutes. Plates were centrifuged at 1,000 x g for 3 min and 100 µL of supernatant was collected, transferred to another 96-well tissue culture treated plate, and absorbance was measured at 412 nm. For CP, 50 µL of 4% normal human sera, 48 µL of GVB++ buffer (0.1 % gelatin, 5 mM Veronal, 145 mM NaCl, 0.025 % NaN_3_, pH 7.3 with 0.15 mM CaCl_2_ and 0.5 mM MgCl_2_, Complement Technology) and 2 µL of inhibitors serially diluted in DMSO were aliquoted into a single well and equilibrated as described. Next, 100 µL of antibody-sensitized sheep red blood cells (Complement Technology) at 5 x10^7^ per well were added to the plate, which was then incubated at 37°C for 1 hour, prior to measurement of absorbance at 412 nm. Data were processed in Graphpad Prism.

### Crystallography and structure determination

6.1 mg/ml C5 (in 20 mM Tris-HCl, 75 mM NaCl, pH 7.35) was mixed at a 1:1 molar ratio with K92_chem_FE. Crystallisation trials were initiated by the hanging drop, vapor-diffusion method at 18 °C by combining C5-K92_chem_FE complex in a 1:1 mixture (v/v) of 0.1 M bicine/Trizma (pH 8.5), 10 % (w/v) PEG 8000, 20 % (v/v) ethylene glycol, 30 mM sodium fluoride, 30 mM sodium bromide, 30 mM sodium iodide. Crystals were flash frozen in liquid nitrogen without additional cryoprotection.

The C5-K92_chem_FE complex crystallised in space group C2 and had unit cell parameters which were isomorphous compared to the C5-K92_bio_ crystals^7^. The C5-K92_chem_FE structure was determined by molecular replacement with Phaser using the C5-K92_bio_ structure as a search model (PDB accession code: 7AD6)^7^. The model was subjected to multiple rounds of manual rebuilding in Coot and refinement in Phenix. The overall geometry in the final structure of the C5-K92_chem_FE complex is good, with 94.4 % of residues in favoured regions of the Ramachandran plot and no outliers. Structure factors and coordinates for the C5-K92_chem_FE complex have been deposited in the PDB (PDB accession code: 7OP0).

### Identifying candidate mutation sites

We visualised the complex crystal structure of K92_chem_FE in complex with complement C5 in PyMOL (version 2.4^55^), with surfaces shown to enable visualisation of cavities and pockets at the interface. Visual inspection revealed a large cavity at the interface of the two complement proteins (chains A and B) with the bovine knob domain. Alanine 12 of K92_chem_FE offered a more direct vector for substitution with a range of non-natural amino acids. We used the Molecular Operating Environment (MOE, version 2019^36^) to screen a small library of commonly used non-natural amino acids. We first prepared the structure using MOE’s standard QuickPrep methodology, which handles protonation and rotamer flipping and other commonly encountered issues with PDB structure files. We then ran MOE’s Residue Scan function, choosing to mutate only Ala 12 to all 20 natural and 50 non-natural amino acids included with the software. This included D variants of most natural amino acids. Both stability difference scores (dStability), based on the overall energy of the complex compared to the unmodified “wild-type” (WT) protein, and affinity difference scores (dAffinity) based on the energies of the knob domain and target protein individually, compared to the bound complex structure, were computed. We noted that MOE frequently inserted rotamers that did not in fact fill the desired pocket but pointed out into solvent. To avoid this, we decided to place virtual atoms (a simple carbohydrate chain) on the outside of the pocket, such that they occupied the same space as the undesired rotamers, thereby producing a steric clash and forcing MOE to choose only side chain orientations that would fill the desired pocket.

This resulted in a ranked list of the 70 candidate amino acids. We selected only those amino acids that were predicted by MOE to improve the complex’s overall stability and the knob domain’s affinity for its target (dAffinity < 0; dStability < 0). This left 26 candidate amino acids, including 8 natural and 18 non-natural amino acids (shown in S8). Rather than further relying on MOE’s energy-based ranking, we took all 18 non-natural amino acids into consideration.

We then manually eliminated a number of candidate side chains due to potentially unstable chemistry as well as difficulties in obtaining the required building blocks to easily synthesize them. Due to their availability, we also chose to synthesise L- and D-ornithine mutants, which were not selected through our MOE analysis. The side chains chosen for synthesis are marked in S9.

### Plasma stability

Rat, mouse and human plasma were collected using lithium heparin as an anti-coagulant. Stability was assessed at a concentration level of 1.25 µg/mL for K57_chem_FE-Palmitoyl, 6.25 µg/mL for K57_chem_FE and 3.75 µg/mL for K8 _chem_FE, over a 24 hour period at room temperature.

A calibration line and suitable quality control samples were prepared and immediately frozen at −80°C, alongside separate spiked samples for the experimental analysis. These separate spiked samples were placed into the freezer after the original calibration line at time intervals of 0.116, 0.25, 0.5, 1, 2, 4, 6 and 24 hours. After a minimum of 1 hour freezing for the final 24 hour spiked samples, the spiked samples were extracted using protein precipitation, alongside the original calibration line.

### Pharmacokinetics

Plasma pharmacokinetics was studied for each peptide in male Sprague-Dawley rats of a body weight between 324 and 425g. Drug was administered intravenously via the tail vein. Doses administered were 10mg/kg for peptides K8_chem_FE, K57_chem_FE and 1mg/kg for K57_chem_FE-Palmitoyl. Blood samples were taken at 7 minutes, 15 minutes, 30 minutes, followed by 1, 2, 4, 8 and 24 hours. Blood was collected into Li Heparin tubes and spun to prepare plasma samples for bioanalysis. Bioanalytical data was analysed on an individual animal basis using Pharsight Phoenix 64 Build 8.1. Non-compartmental analysis was conducted and mean pharmacokinetic parameters for each drug calculated.

### Bioanalysis sample preparation

Calibration samples were prepared in lithium heparin plasma from DMSO stock solutions. For K8 _chem_FE this was over a linear range of 0.1 to 250µg/mL; for K57 _chem_FE from 0.05 to 125µg/mL and for K57 _chem_FE-Palmitoyl over a linear range of 0.025 to 12.5 µg/mL.

20 µL of plasma samples were distributed in a 96 well microtitre plate. To each sample, 25 µL of internal standard at a concentration of 25 ng/mL in methanol was added and vortex mixed. The internal standards were Metoprolol, For K57_chem_FE and K8_chem_FE, and Propranolol for K57_chem_FE-Palmitoyl. 100 µL of methanol:formic acid (100:0.2) was added to each sample to precipitate the plasma proteins. The plate was then shaken for 2 minutes and then centrifuged at 2500g for 8 minutes at 5 °C. Finally, 50 µL of supernatant was transferred to an empty 96-well analytical plate using a Hamilton liquid handler and diluted with 100 µL of water:formic acid (100:0.2).

### Bioanalysis (LCMS/MS)

The analytical plate was loaded into a Waters Acquity iClass UPLC systemand 3-15 μL of the extracted samples injected onto the LC–MS/MS system. The cyclic peptides were separated on a Waters Acquity UPLC column (BEH C18 1.7 μm 2.1 mm × 50 mm). The UPLC column was equilibrated with the initial mobile phase conditions at a flow rate of 0.4 mL/min and the analytes were separated using a water, acetonitrile and formic acid gradient described below. Solvent A was 100 % water 0.1 % formic acid and solvent B 100 % acetonitrile.

For K8_chem_FE: After a 0.2 minute hold, a gradient from10 % acetonitrile (B) increasing to 30 % solvent B over 3 minutes was performed. For K57_chem_FE: After a 0.2 minute hold, a gradient from 0% acetonitrile (B) increasing to 25% solvent B over 3 minutes was performed. For K57 _chem_FE-Palmitoyl: After a 0.2 minute hold, a gradient from 20 % acetonitrile (B) increasing to 50% solvent B over 3 minutes was performed. For all three peptides from 3.0 to 3.4 min solvent B increased to 95 %.

This was held at 95 % from 3.4 to 4.4 minutes then back to 0 % solvent B at 4.5 min and held until 5.0 min, in preparation for the next sample injection. The retention time for the analytical and reference internal standard was 2.13 minutes for K8_chem_FE, 2.45 minutes for K57 _chem_FE, and 2.25 minutes for K57 _chem_FE-Palmitoyl. The internal standards eluted at 2.05 minutes and 1.31 minutes for Metoprolol and Propranolol, respectively.

The analytes were detected by turbo ionspray ionization and MS/MS detection using an AB Sciex 6500+ MS/MS triple quadrupole mass spectrometer. The analytical column eluate was delivered into the source operated with a positive IonSpray voltage and operated using multiple reaction monitoring (MRM) mode.

MRM was carried out by setting at a precursor ion (m/z) and fragment ion showing a satisfactory detection sensitivity as a product ion. For the intact peptides, MRM transitions were determined in positive ion mode, via multiply charged precursor ions (m/z) and singly charged fragment ions. These corresponded to m/z ratios of: [M + 6]+ 845.0 >973.7 for K8_chem_FE., [M + 5]+ 821.1>985.5 for K57 _chem_FE and +5 [M + 4]+ 1165.4>730.2 for K57 _chem_FE-Palmitoyl. The MRM transitions for the internal standards were [M + 1]+ 268.2>159.1 for metroprolol and [M + 1]+ 260.1>155.0 for propranolol. The ion chromatograms were quantified by reference to standard curves spiked into fresh control lithium heparin plasma and by plotting the analyte:internal standard peak area ratio of the calibration standards vs. the concentration of peptides. The concentration of peptide in the *in vivo* study, QC and stability samples was determined by interpolation of the peak area ratios from the calibration curve.

## Acknowledgements

We would like to thank Dr Michele Farris, from Peptide Protein Research, Callum Lord Mears, from Covance Ltd, as well as Anthony Veltri and Judith van Asperen, from UCB, for their assistance in coordinating this study.

## Author contributions

The study was designed and supervised by A.M., J.v.d.E. and A.D.G.L. Peptide synthesis was performed by J.H. and R.J.B. Protein purification and SPR was performed by A.M. LC/MS on capped and reduced peptides was by A.H. The complement activation ELISA were performed by A.M., C.J., Y.T. and D.V., with haemolysis assays by Y.T. and D.V. The C3d ELISA counter screen was performed by T.W. and M.L. Crystallography was performed by A.M., J.R.B. and J.v.d.E. The rational design of K92_chem_FE mutants was performed by S.K., R.T., M.D.S., R.J.F. and J.S. The *in vivo* study was designed by K.B., with bioanalysis and *in vitro* plasma stability performed by G.B. and K.S. The manuscript was written by A.M., with all authors providing critical input and review.

## Supplementary Material

**S1.**
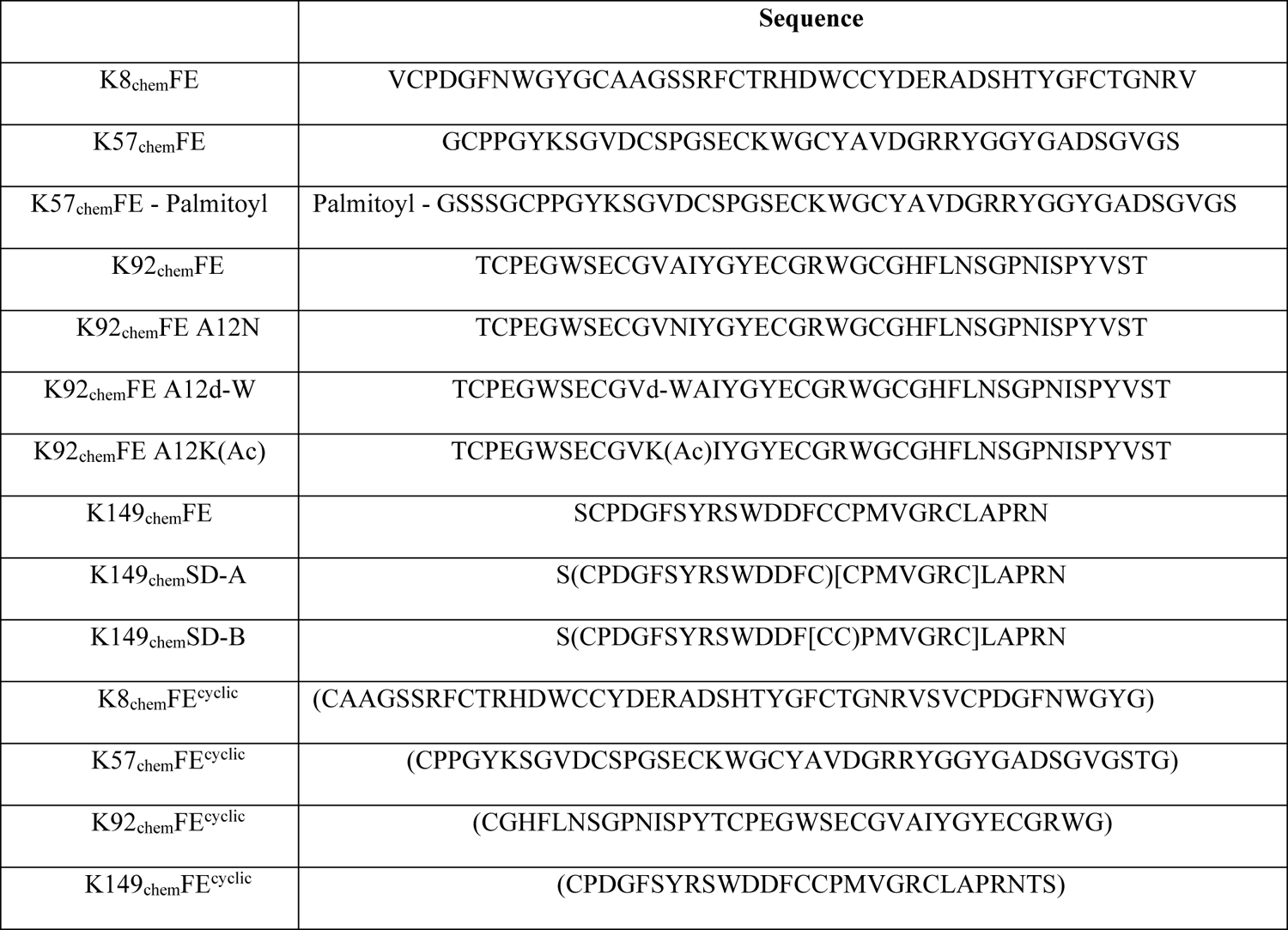
Knob domain sequences that were chemically synthesised in this study

**S2.**
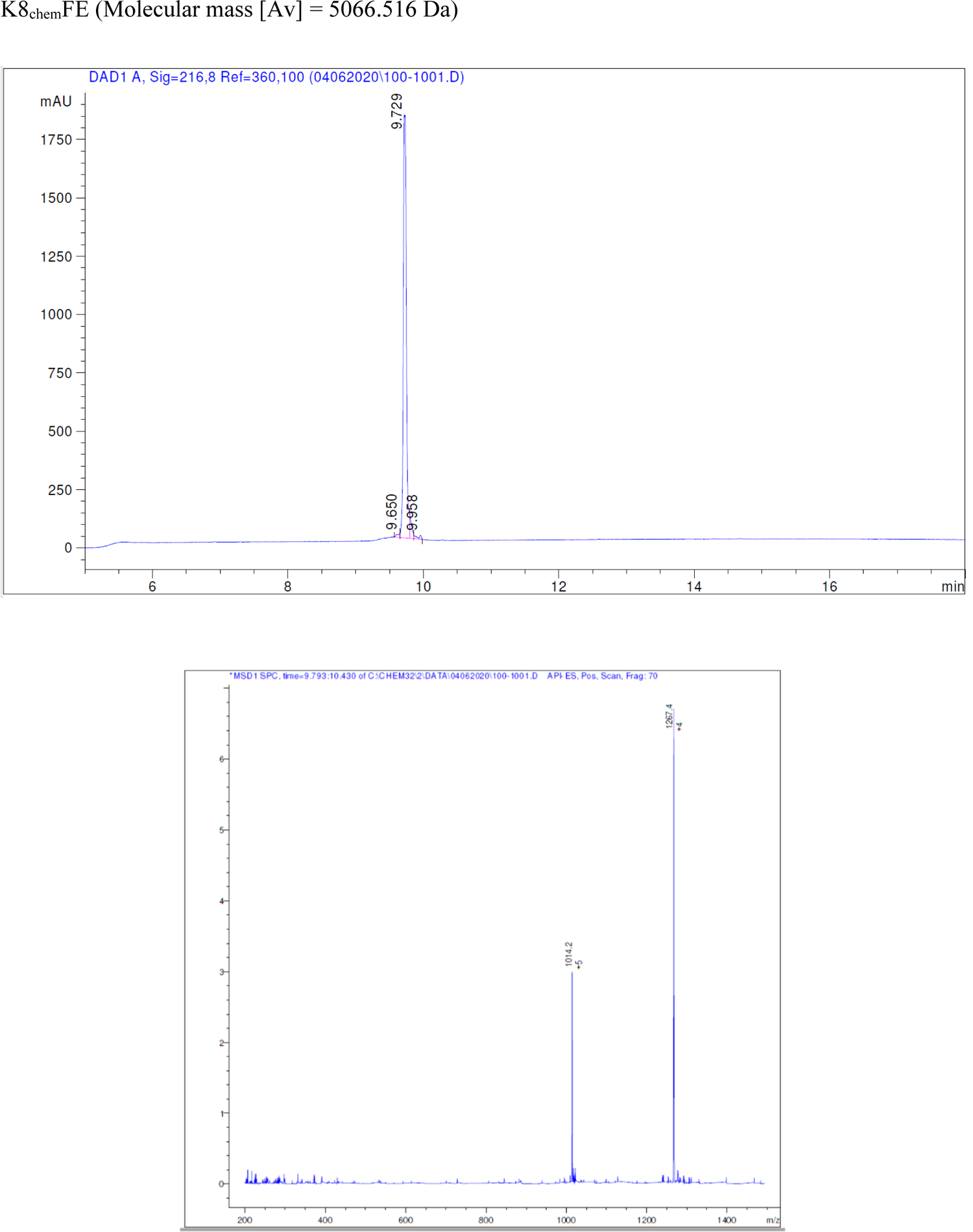
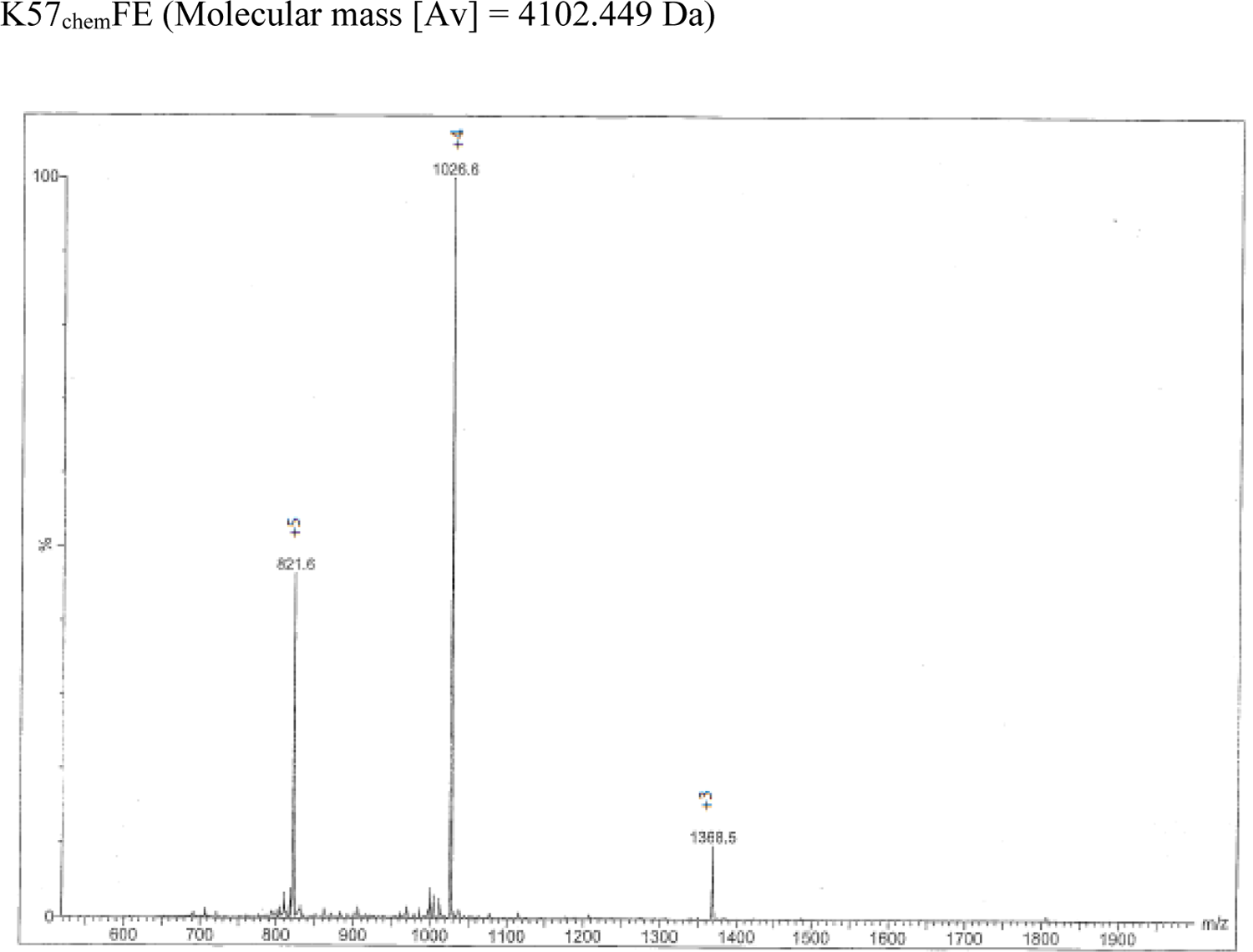
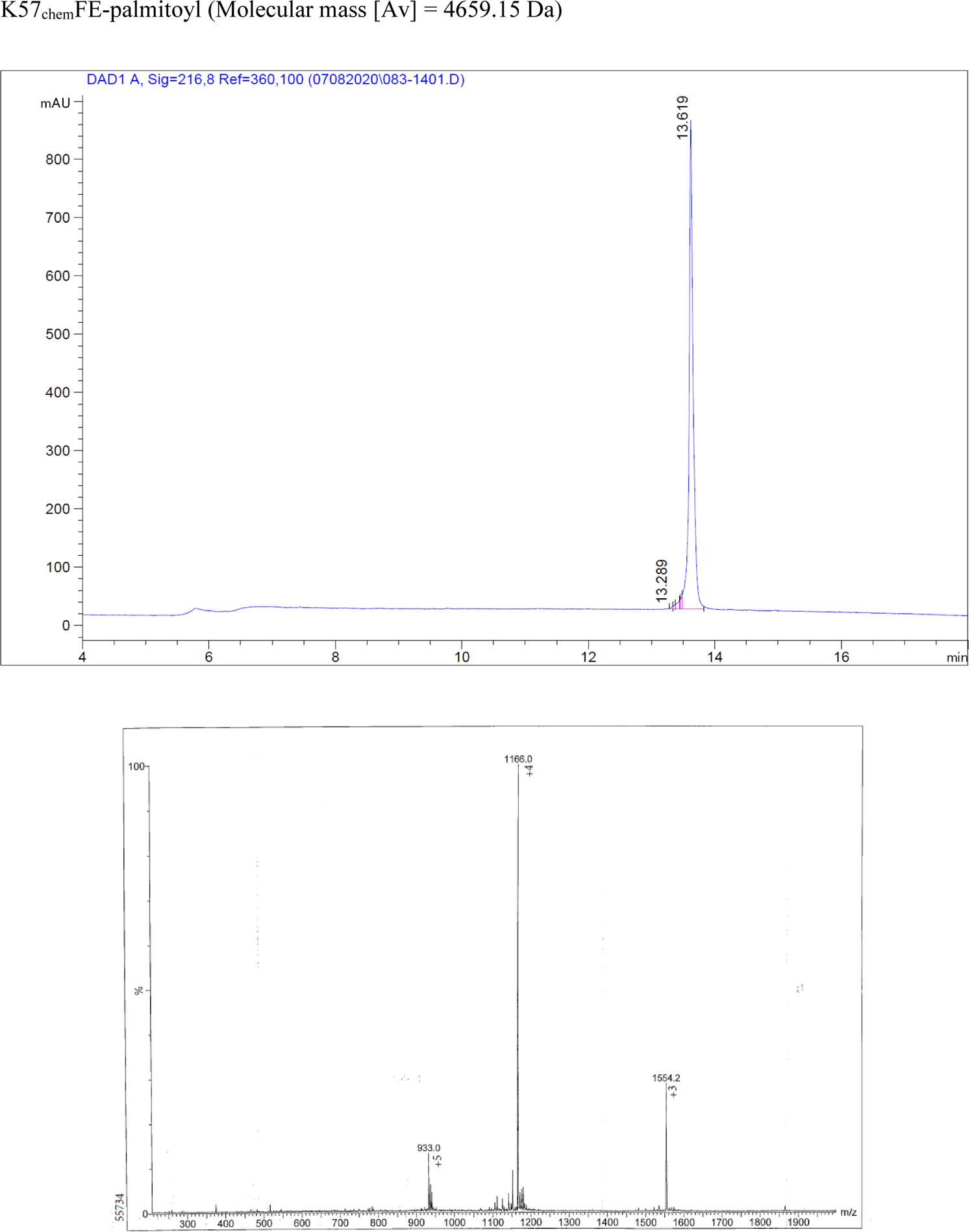
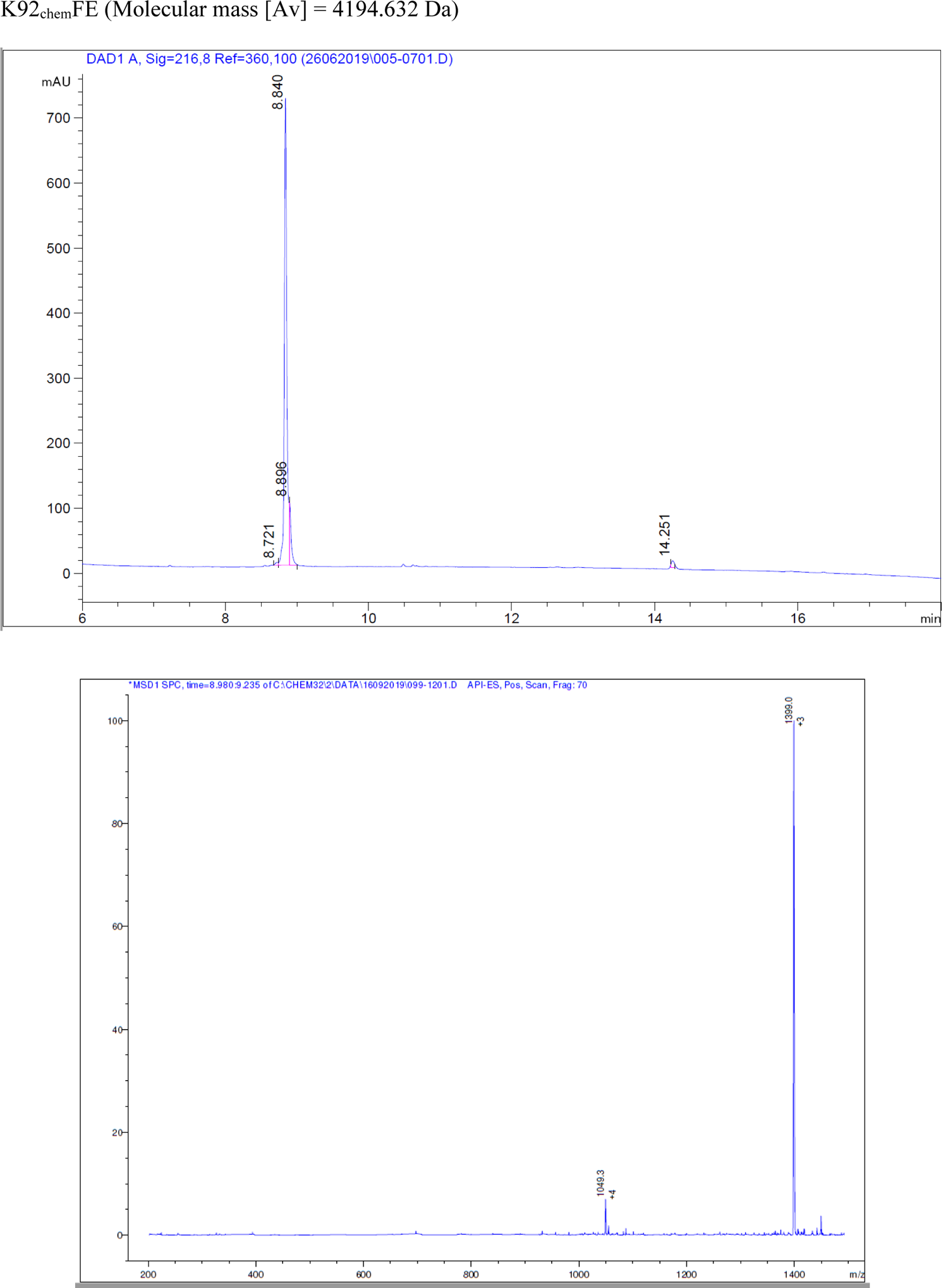
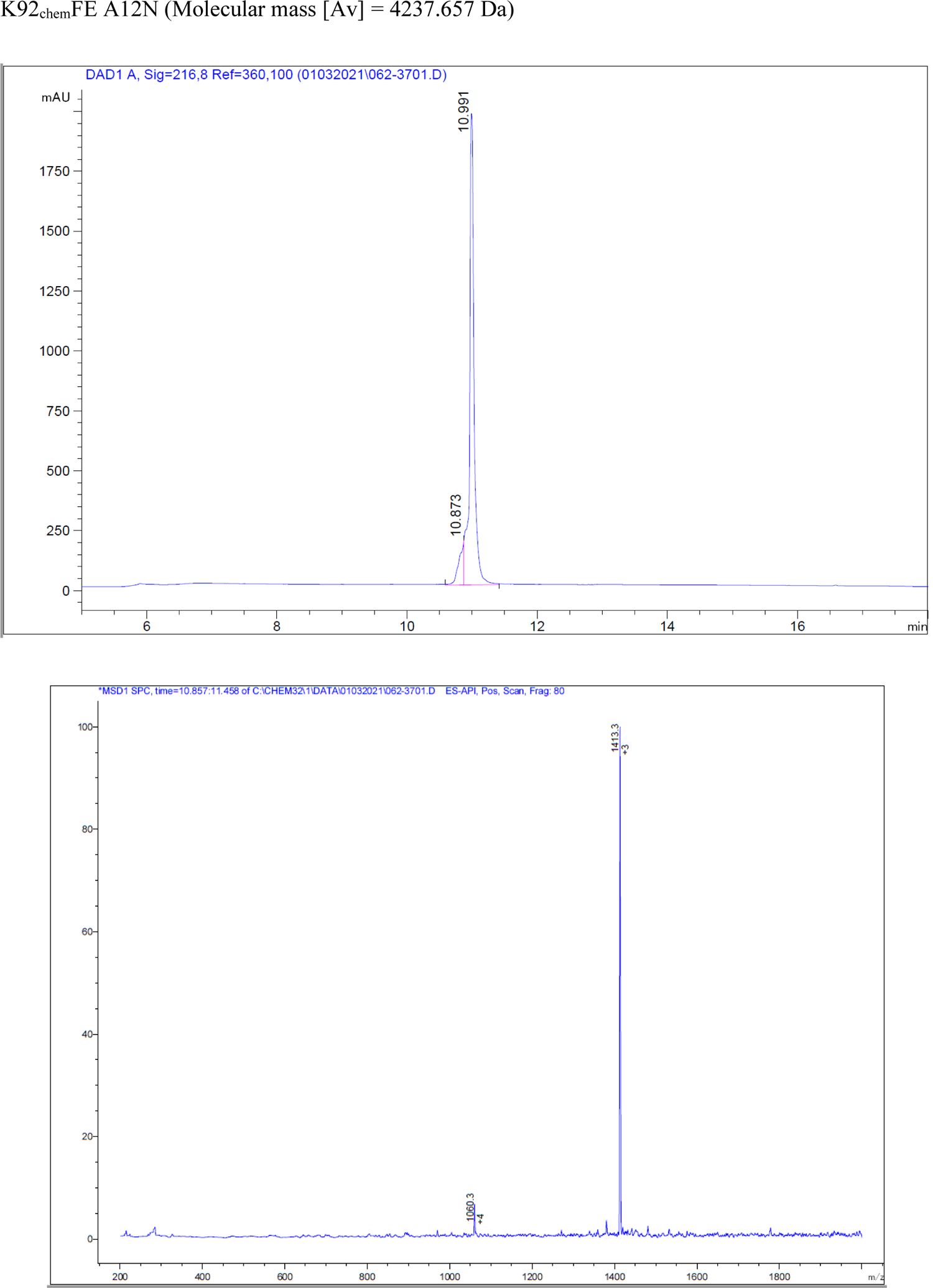
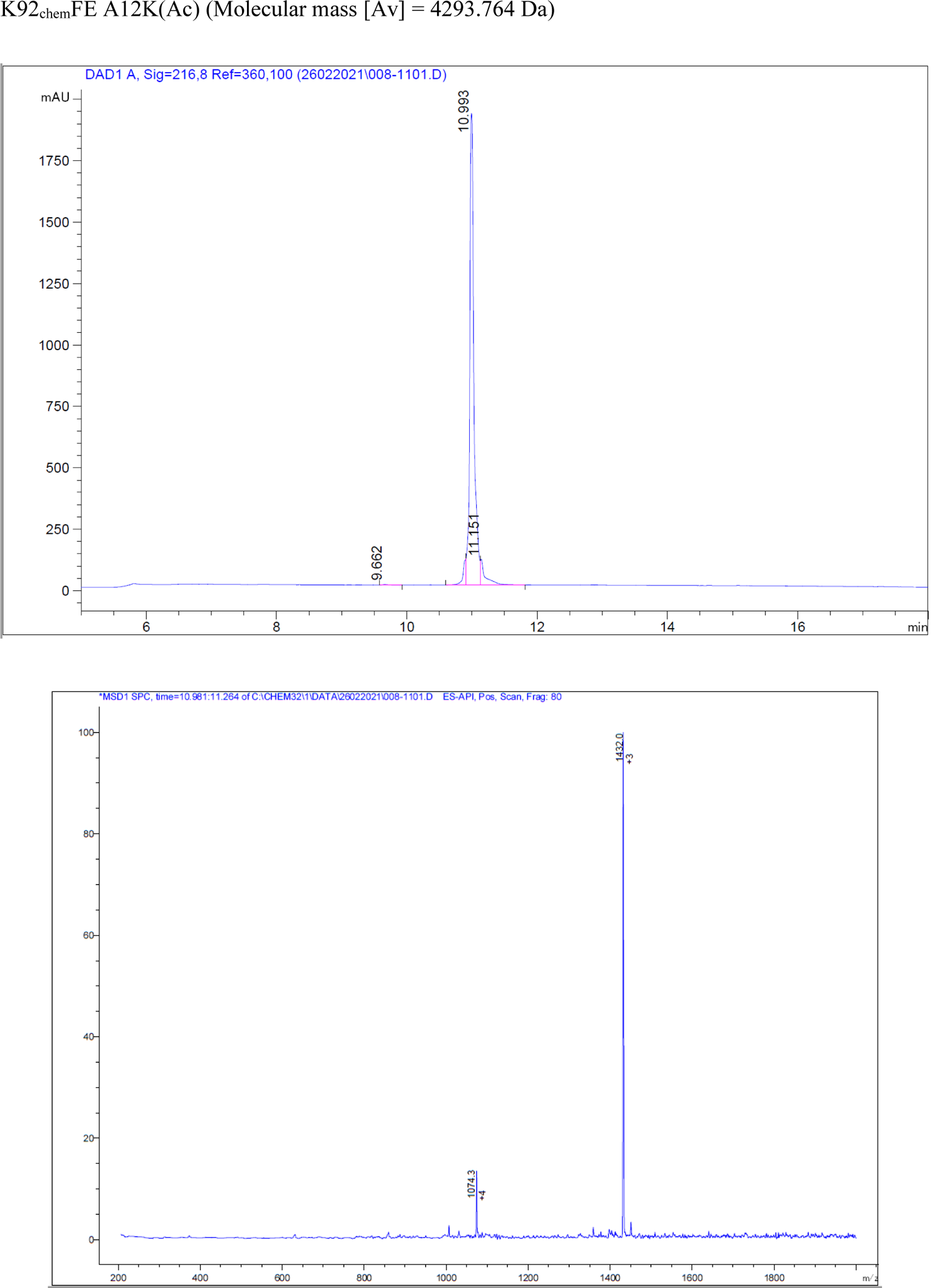
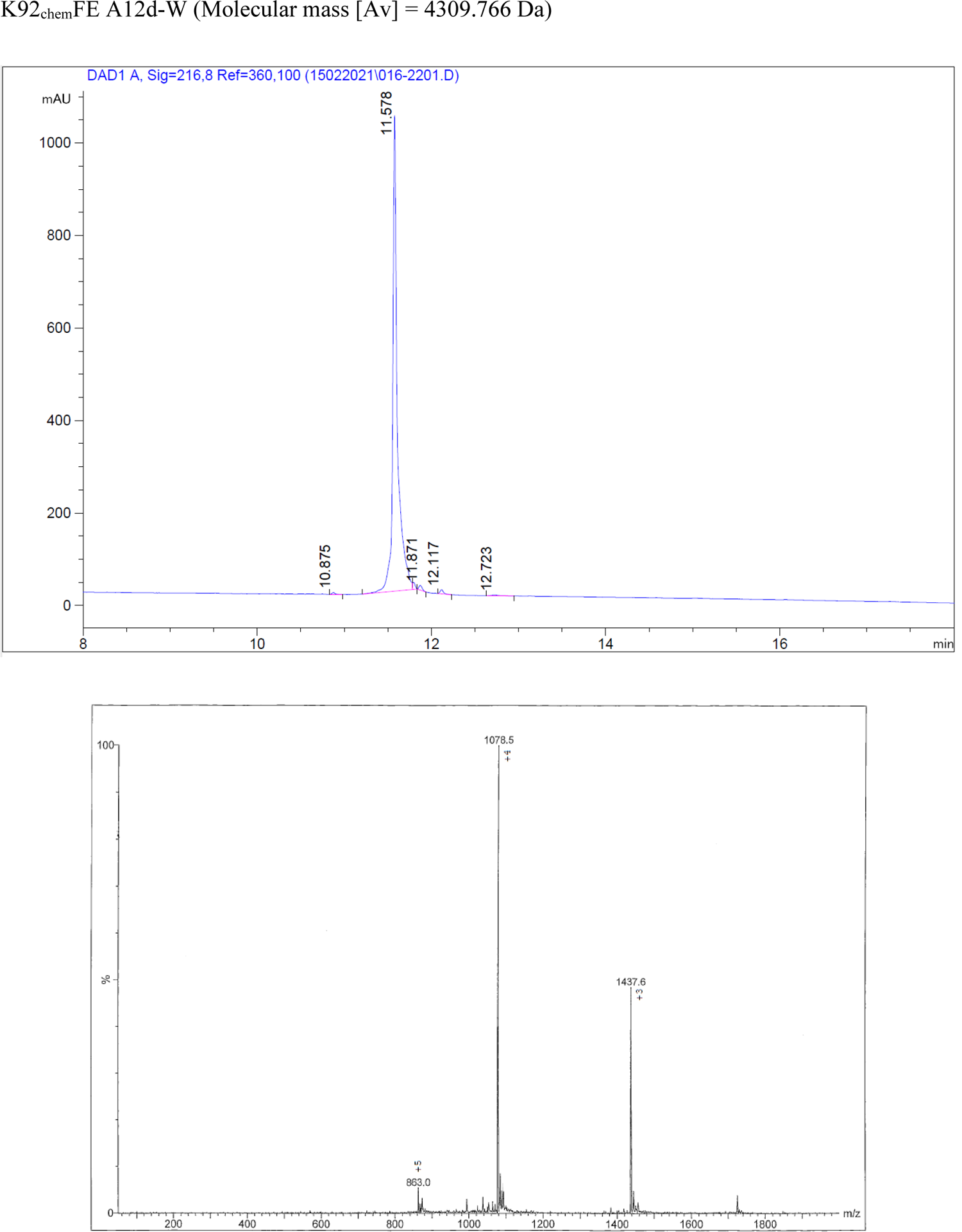
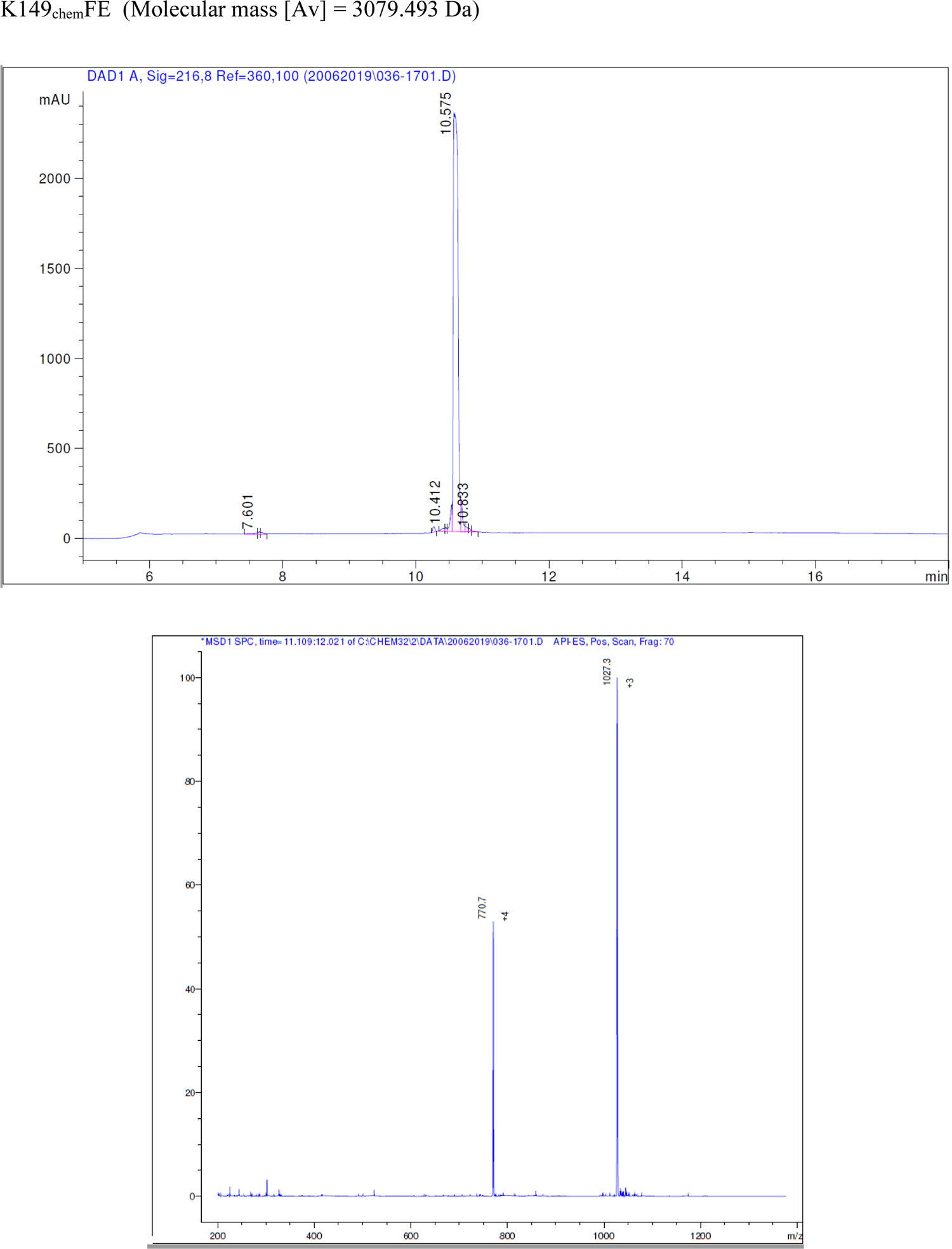
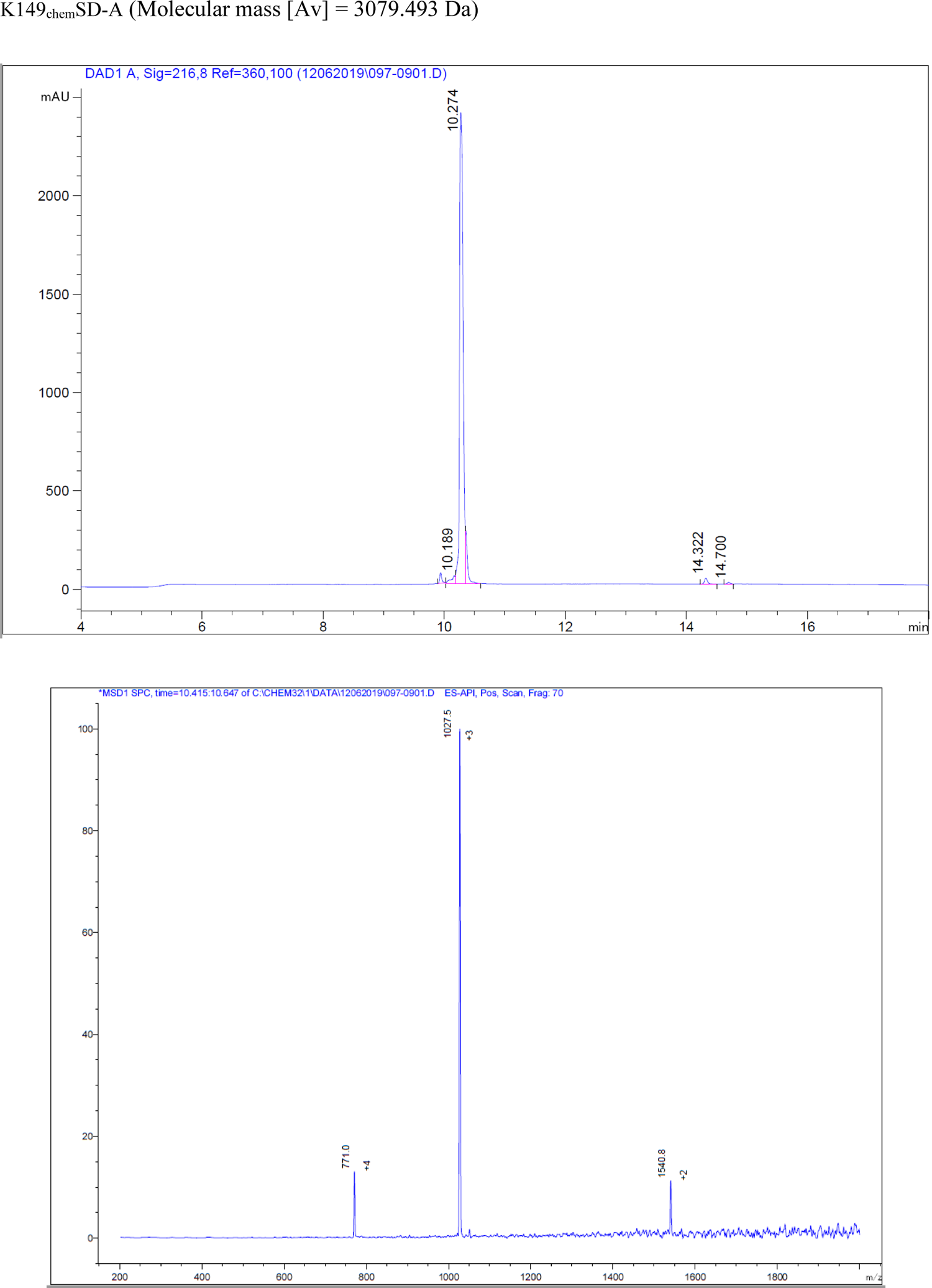
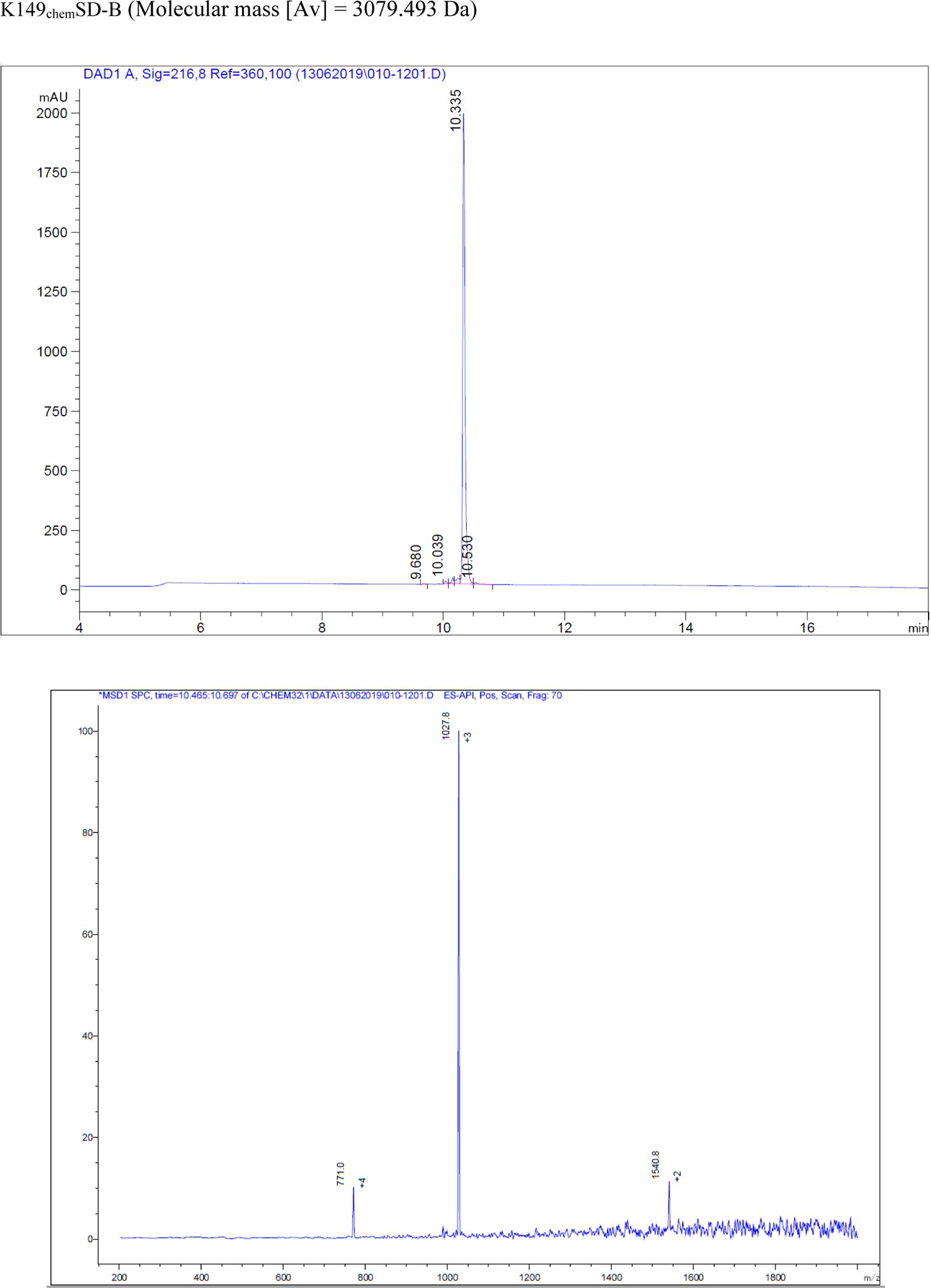
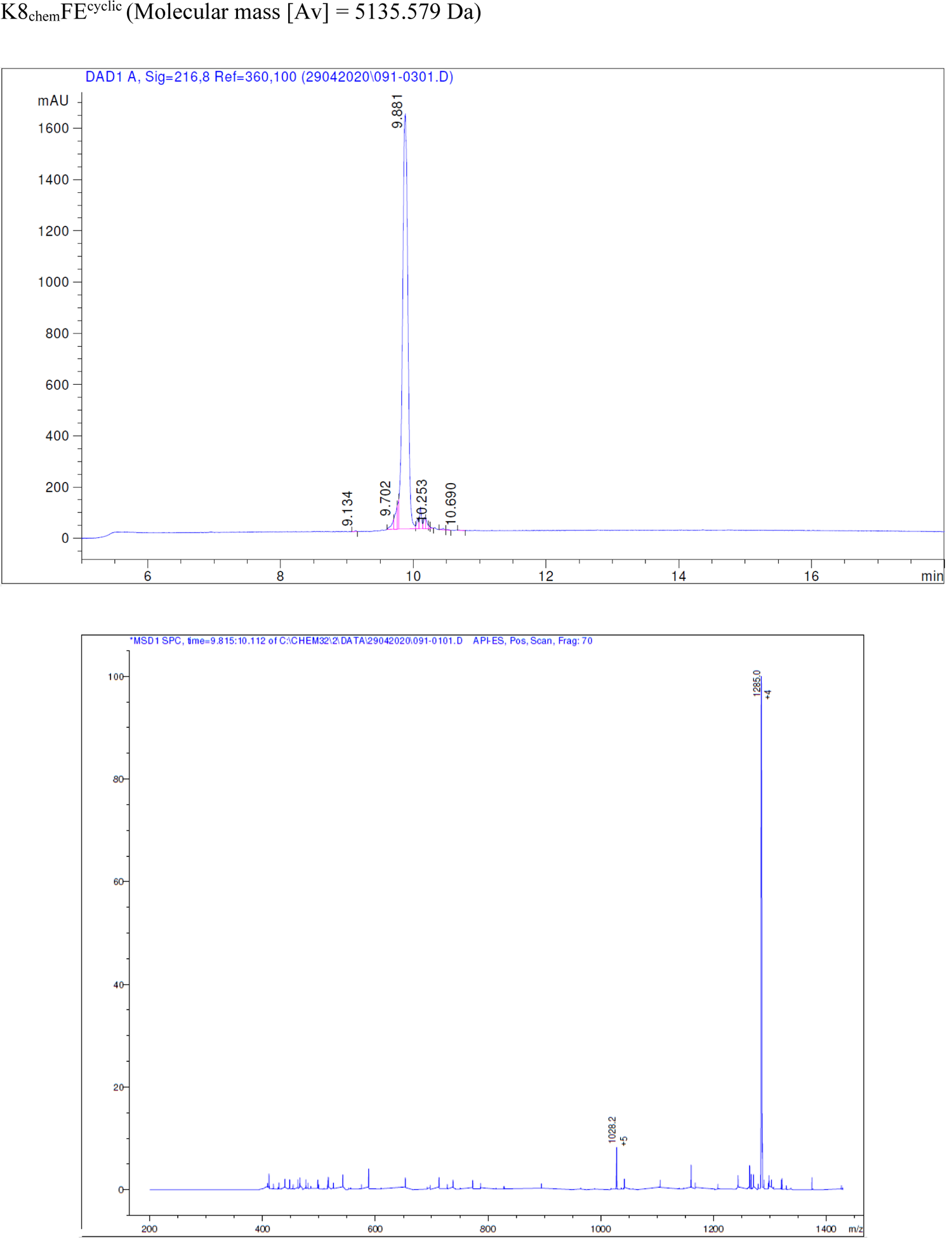
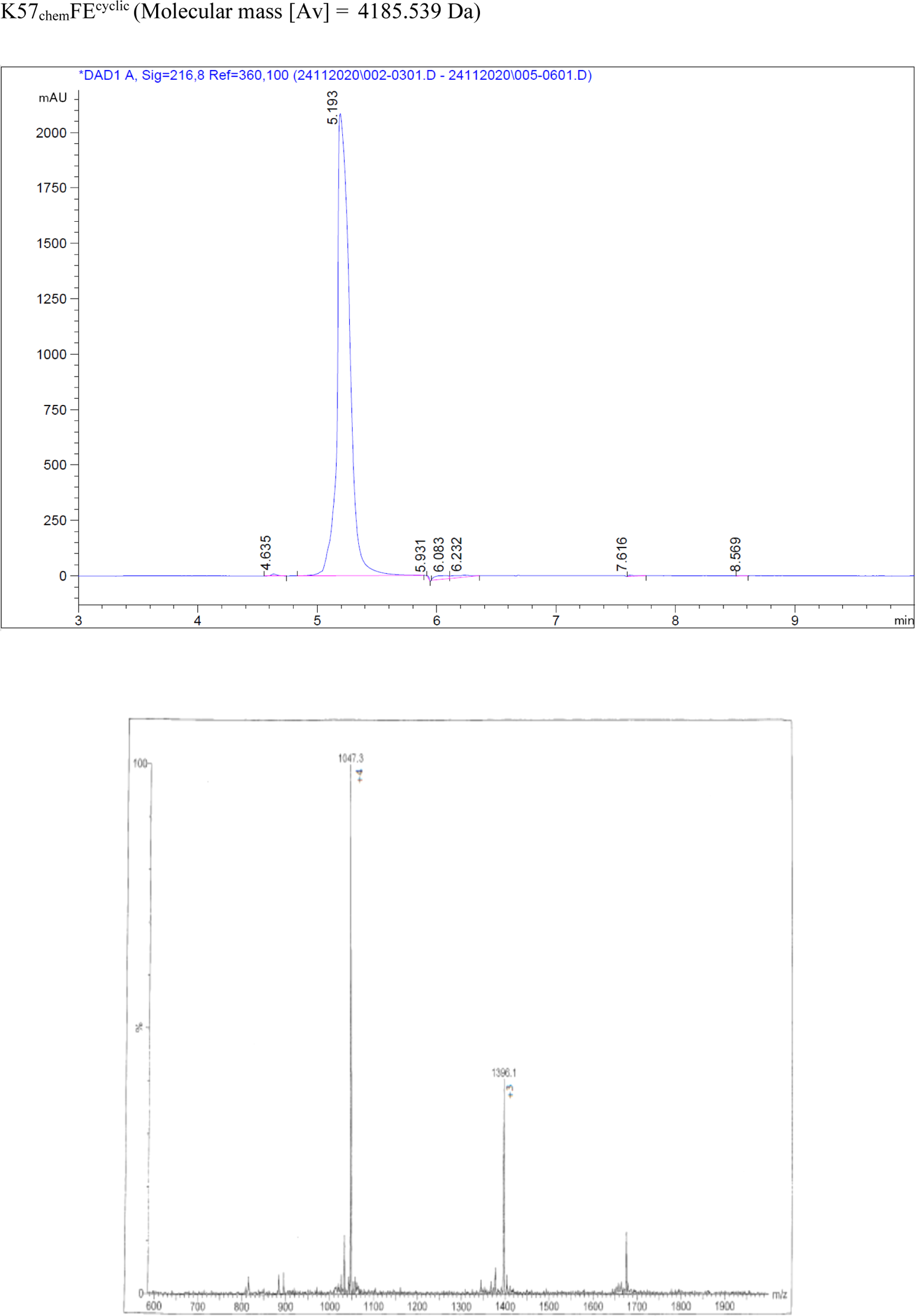
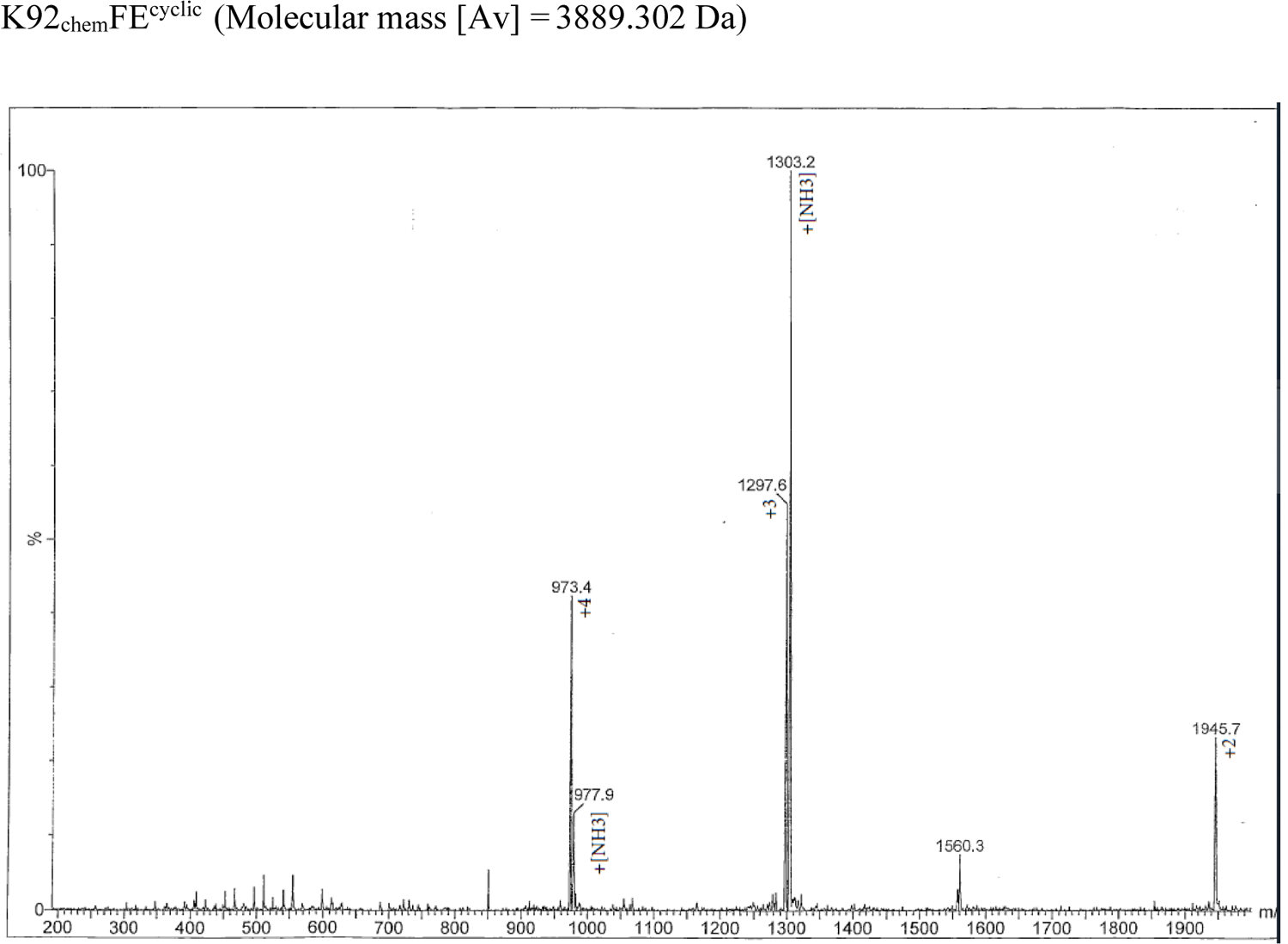
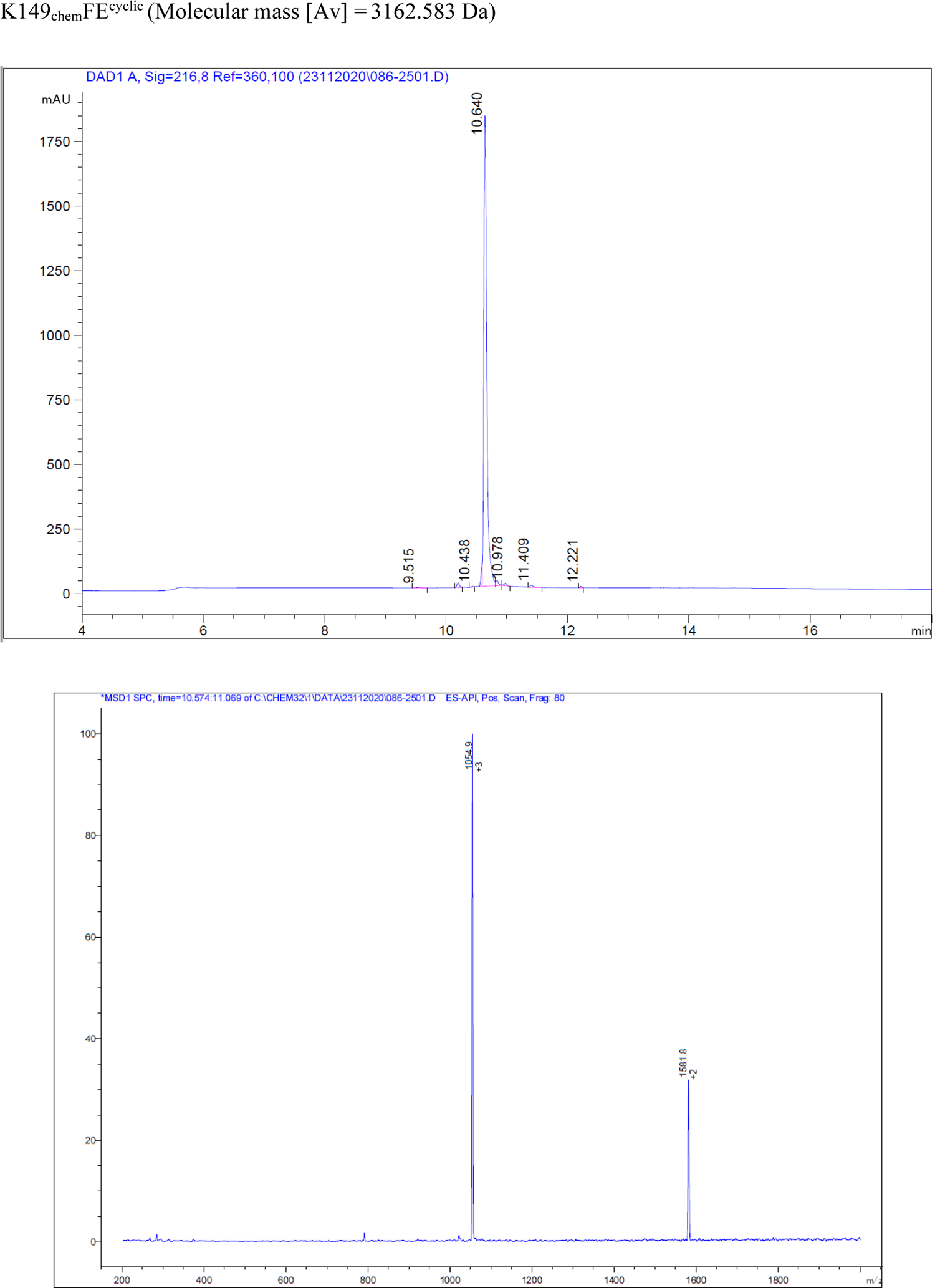
Knob domain QC: HPLC chromatograms and MS spectra

**S3.**
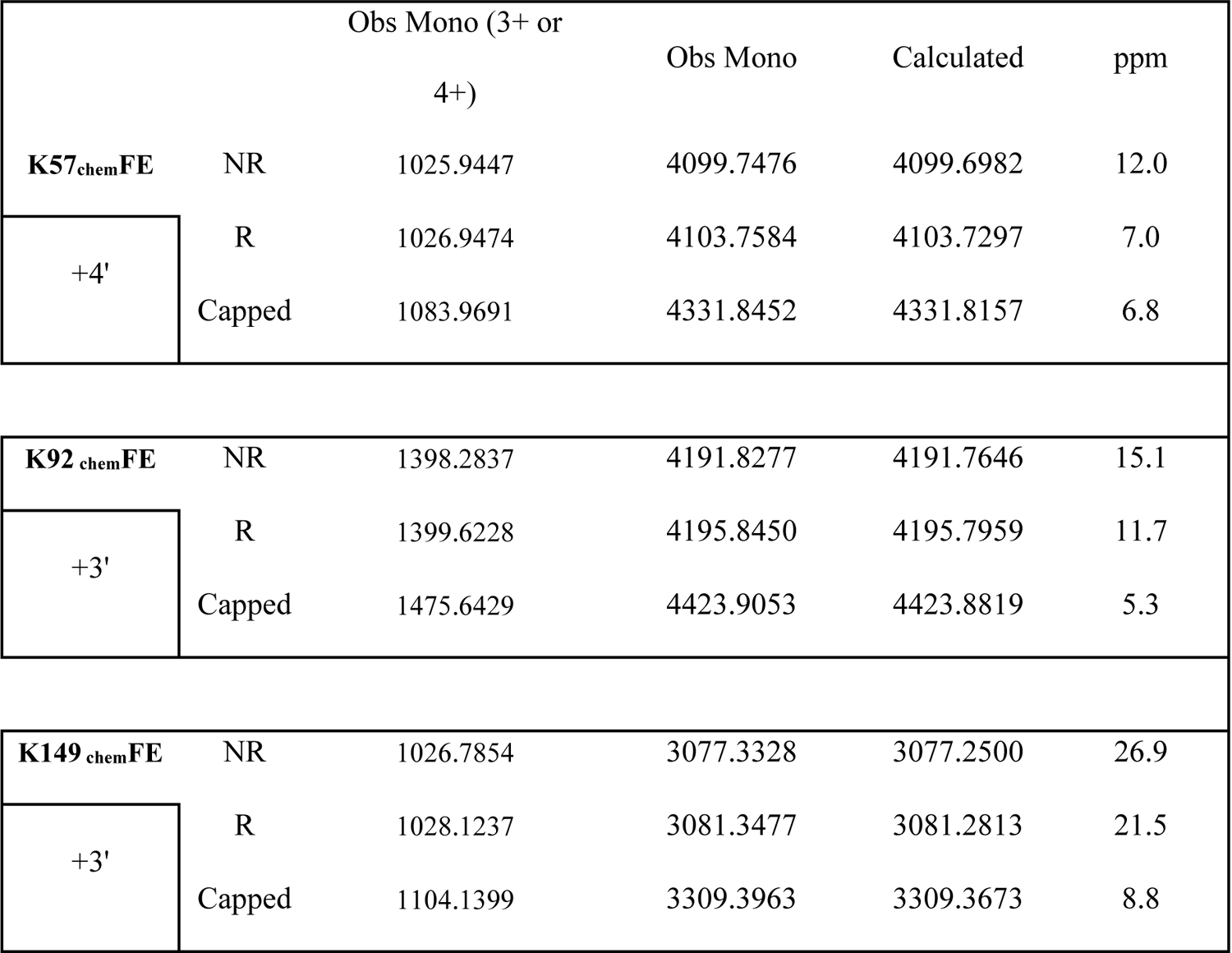
Mass Spectrometry to confirm capping of knob domain peptides. To confirm capping, reduced (R) and non-reduced (NR) knob domains were analysed alongside knob domains which had been reduced and capped with IAM. For all samples, the observed monisotopic masses (Obs Mono) show increases in molecular weight which corresponding to the reduction of disulphide bonds, in the R samples, and addition of IAM, in the capped samples, confirming that the cysteines have been universally capped and are not participating in disulphide bonds, with reducing and capping resulting in a mass change of +58 per cysteine.

**S4.**
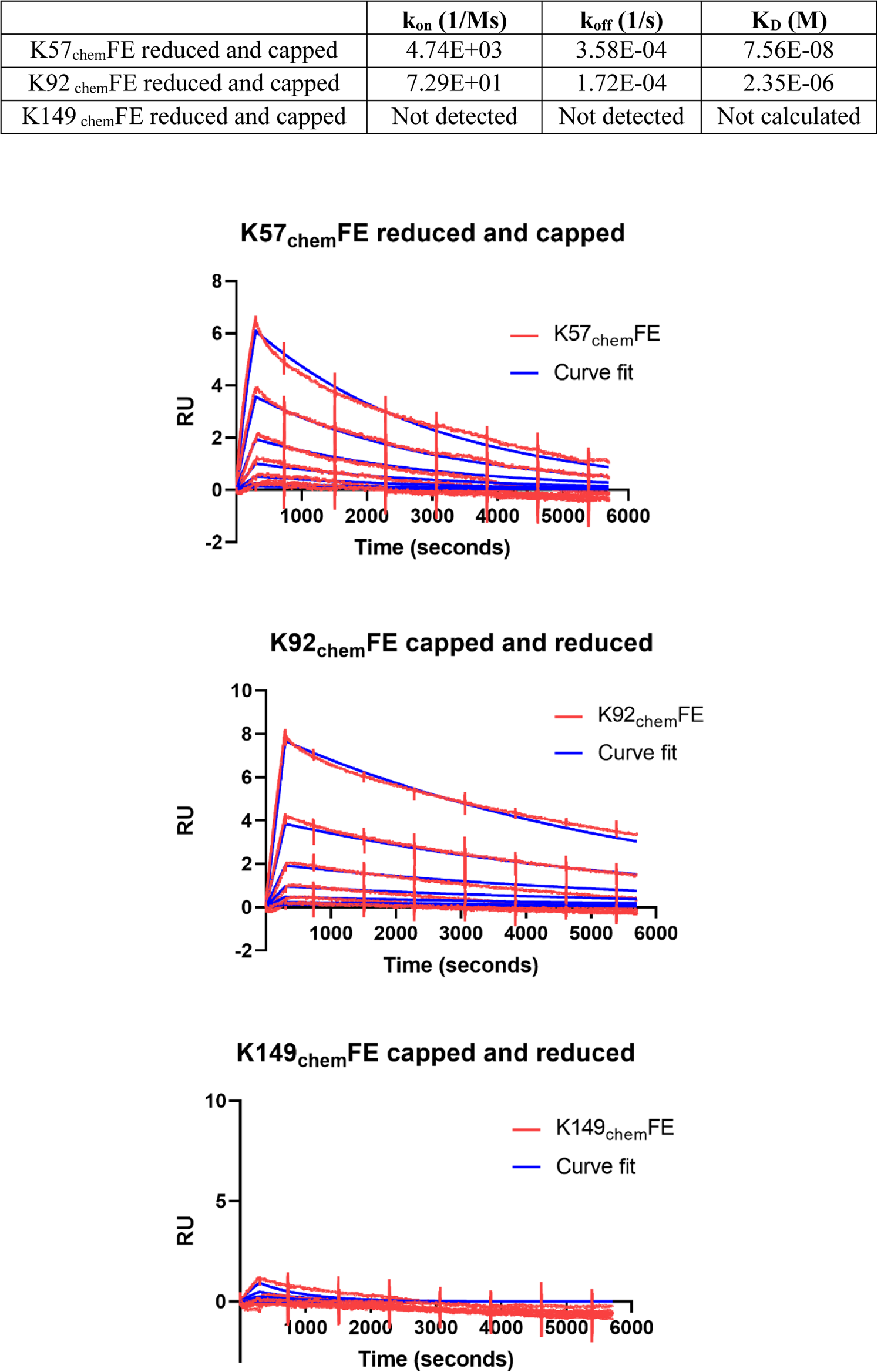
SPR on reduced and capped knob domains

**S5.**
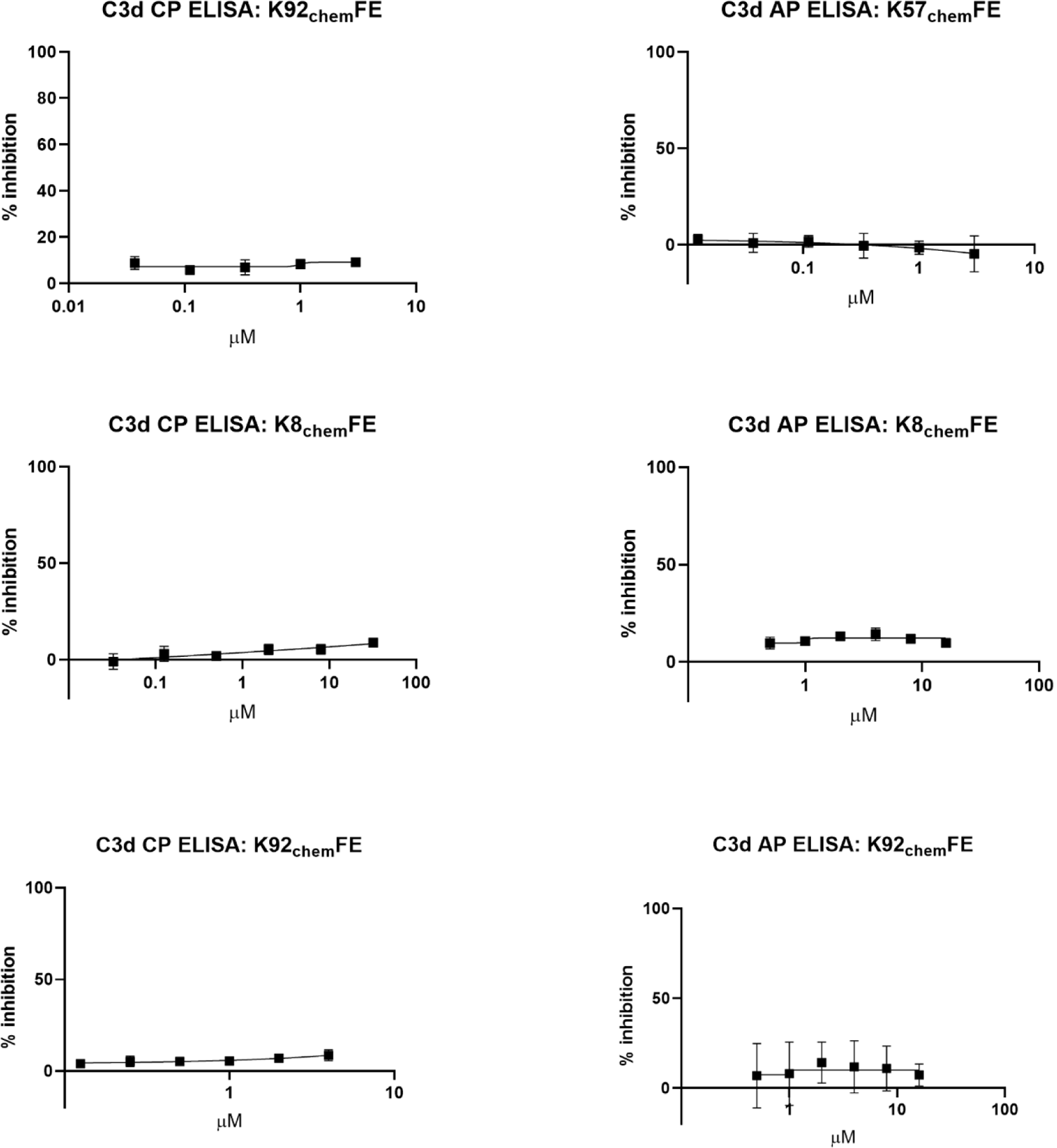
C3d Activation ELISA counterscreen, data from *n=3* experiments

**S6.**
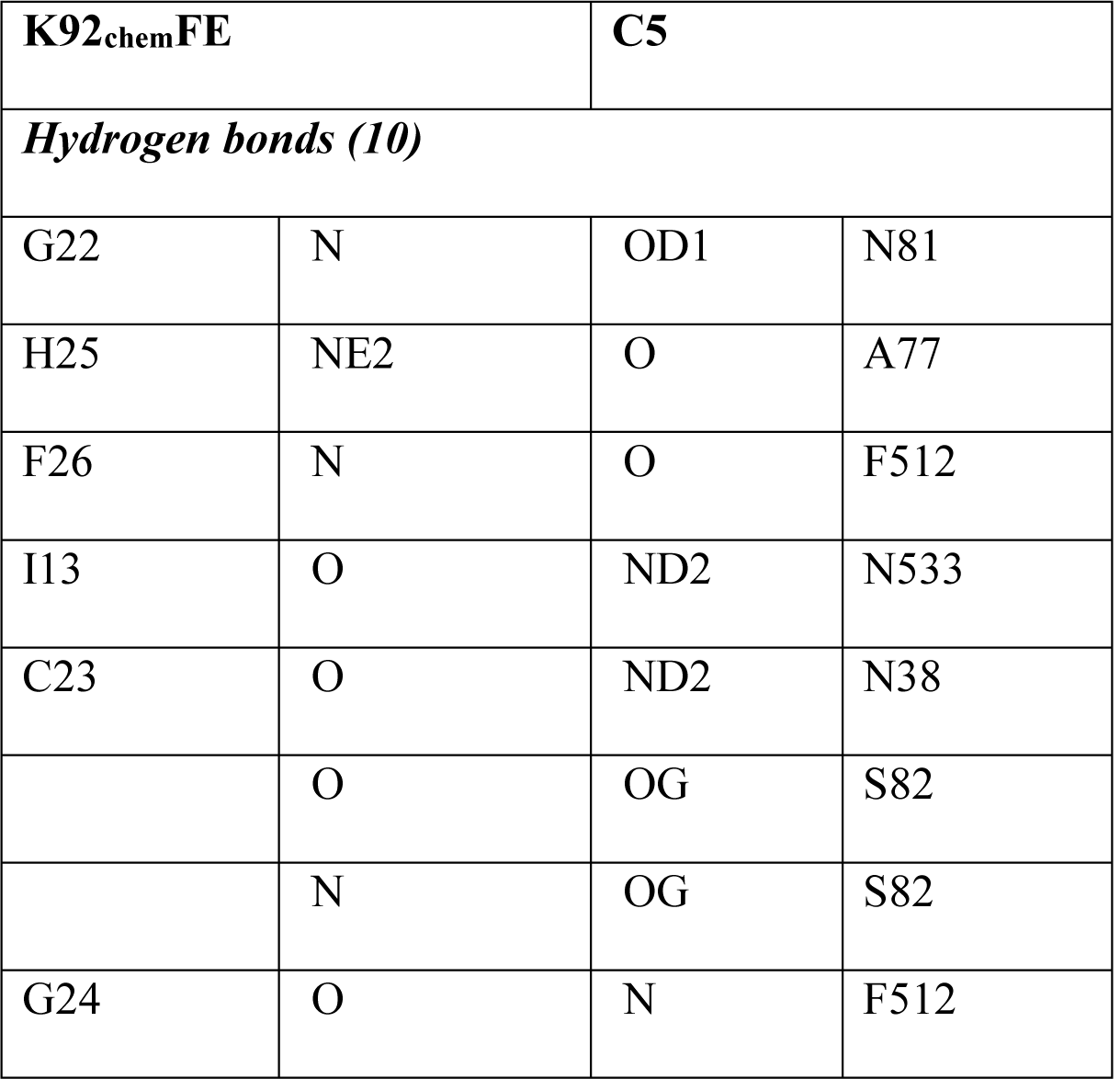
Table of hydrogen bonds between K92chemFE and C5. Hydrogen bonding interactions as defined by the PDBePiSA macromolecular interfaces tool.

**S7.**
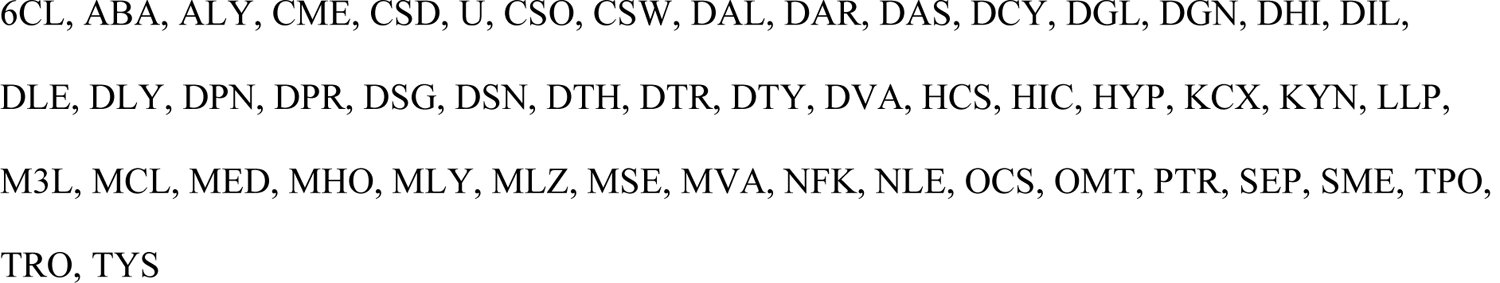
Full list of symbols for non-natural amino acids scanned in MOE (50 total):

**S8.**
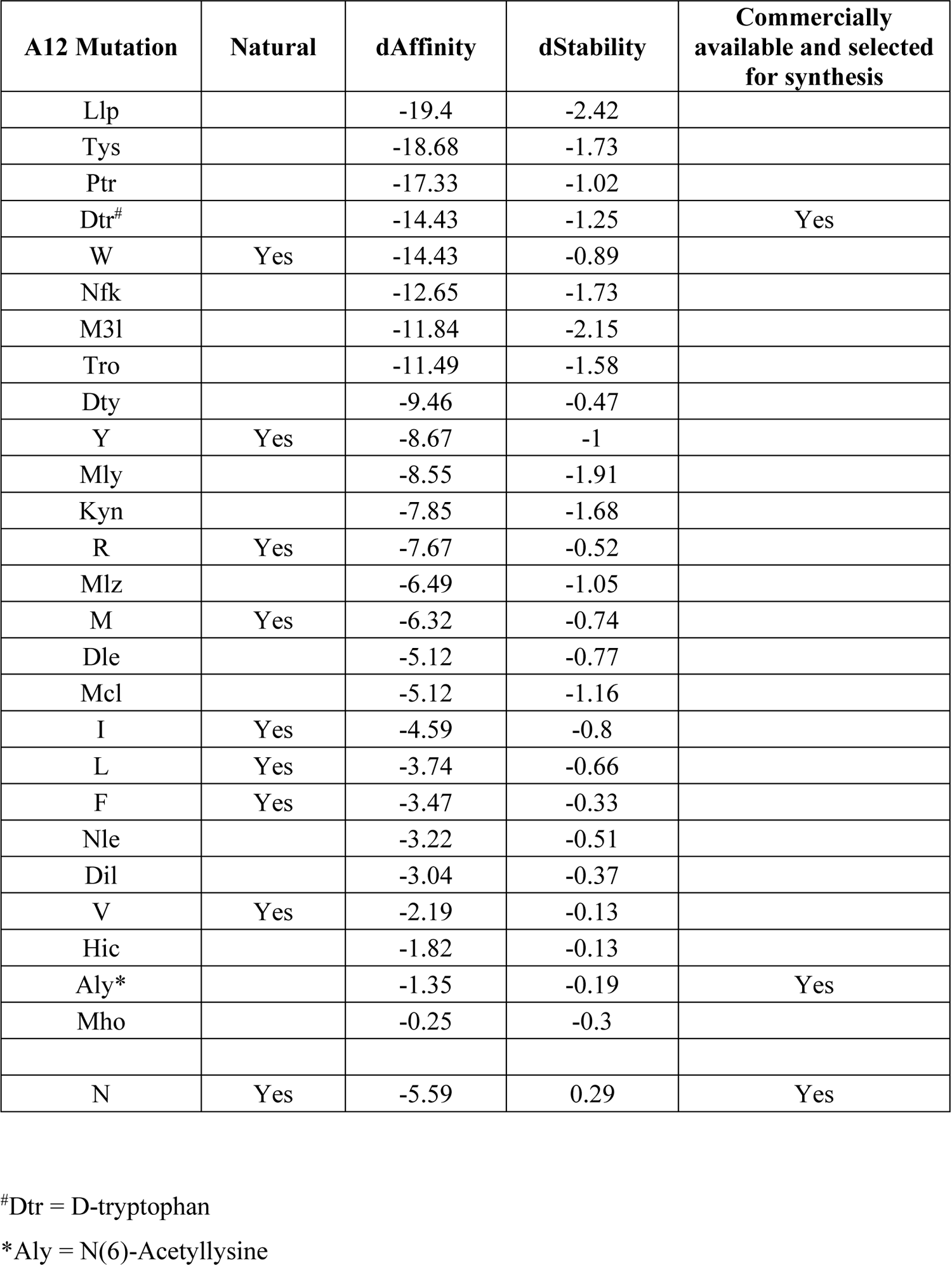
Amino acids suggested by MOE, for substitution of K92’s residue 12, after applying the filters: dAffinity < 0; dStability < 0.

**S9.**
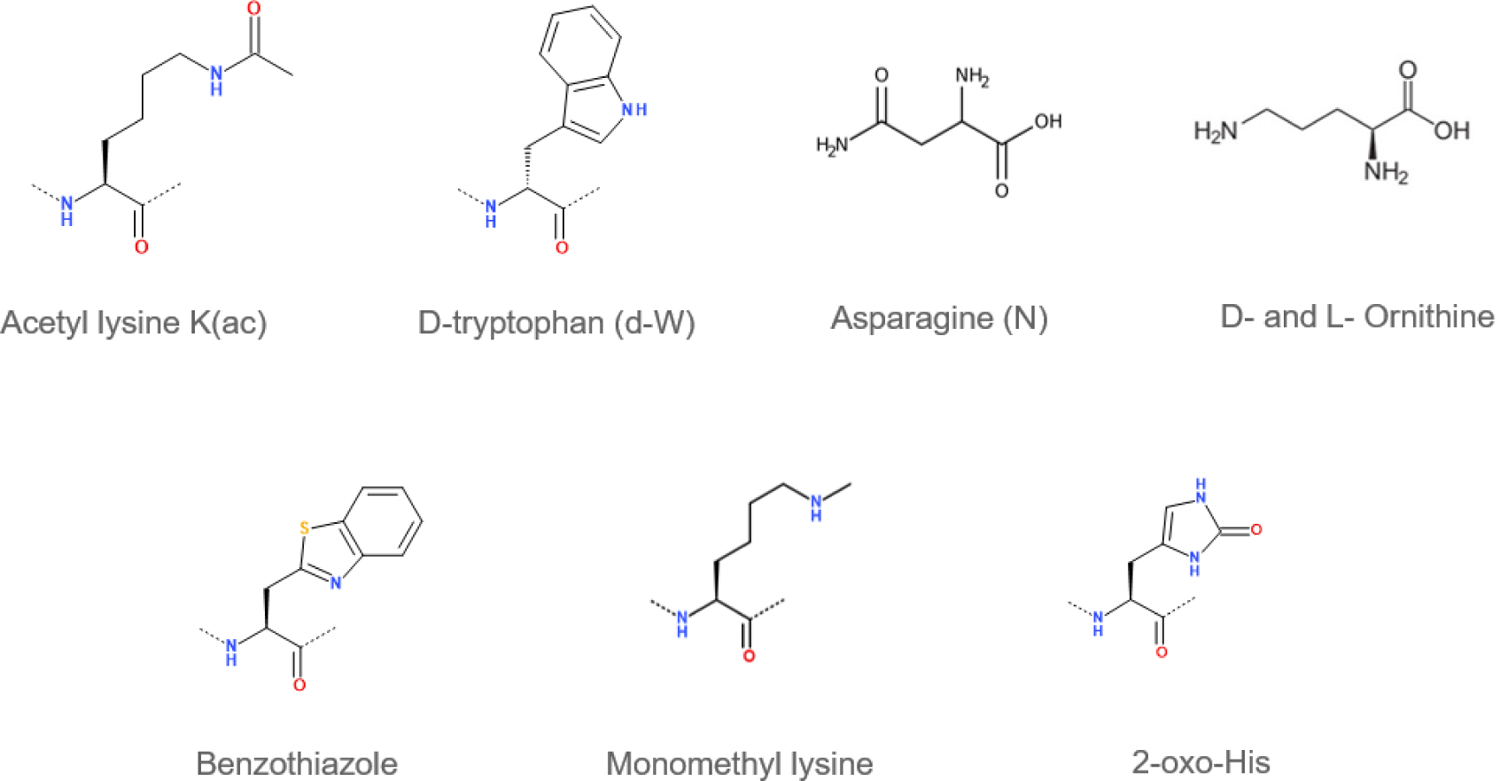
Amino acids exemplified as K92**_chem_**FE Alanine 12 mutations

**S10.**
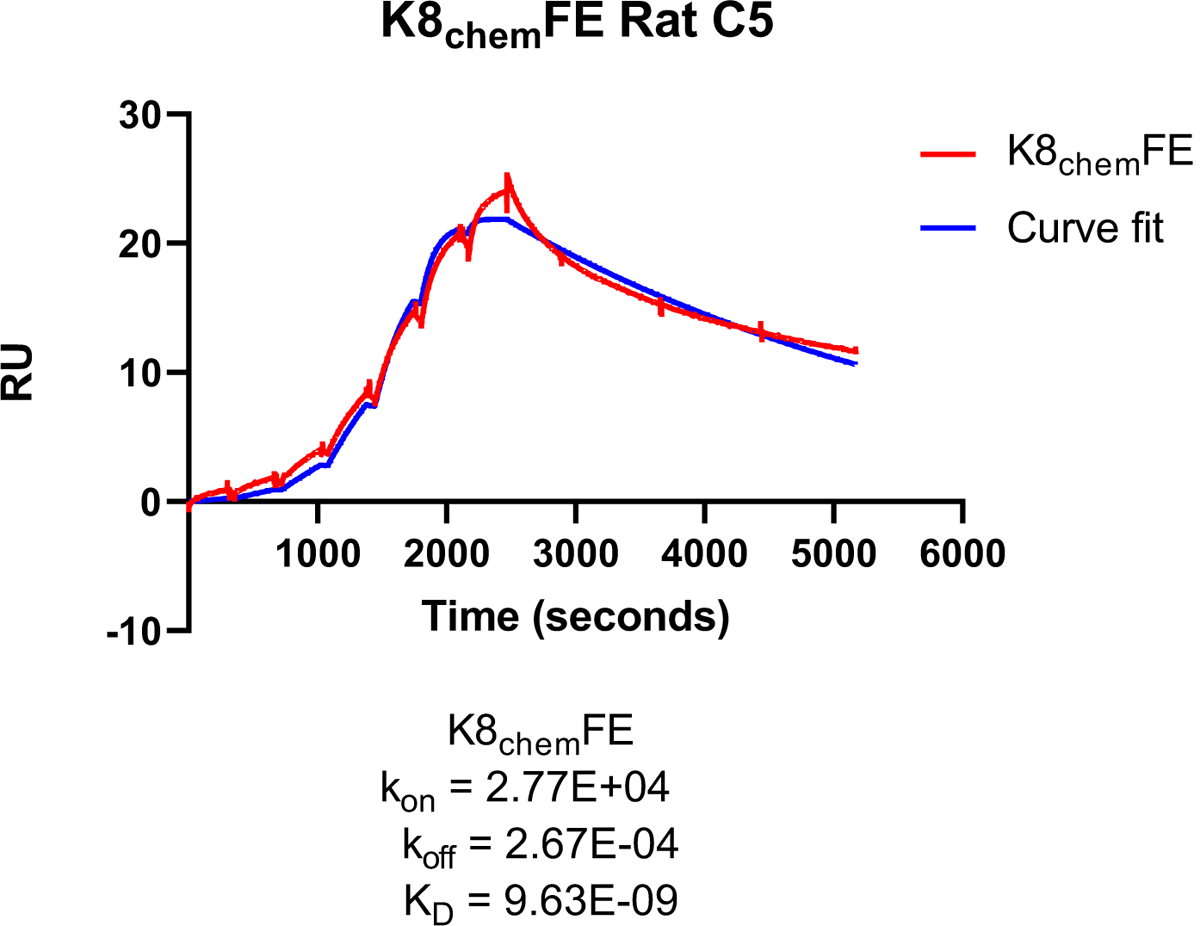
Binding of K8_chem_FE to rat C5 by SPR single-cycle kinetics.

**S11.**
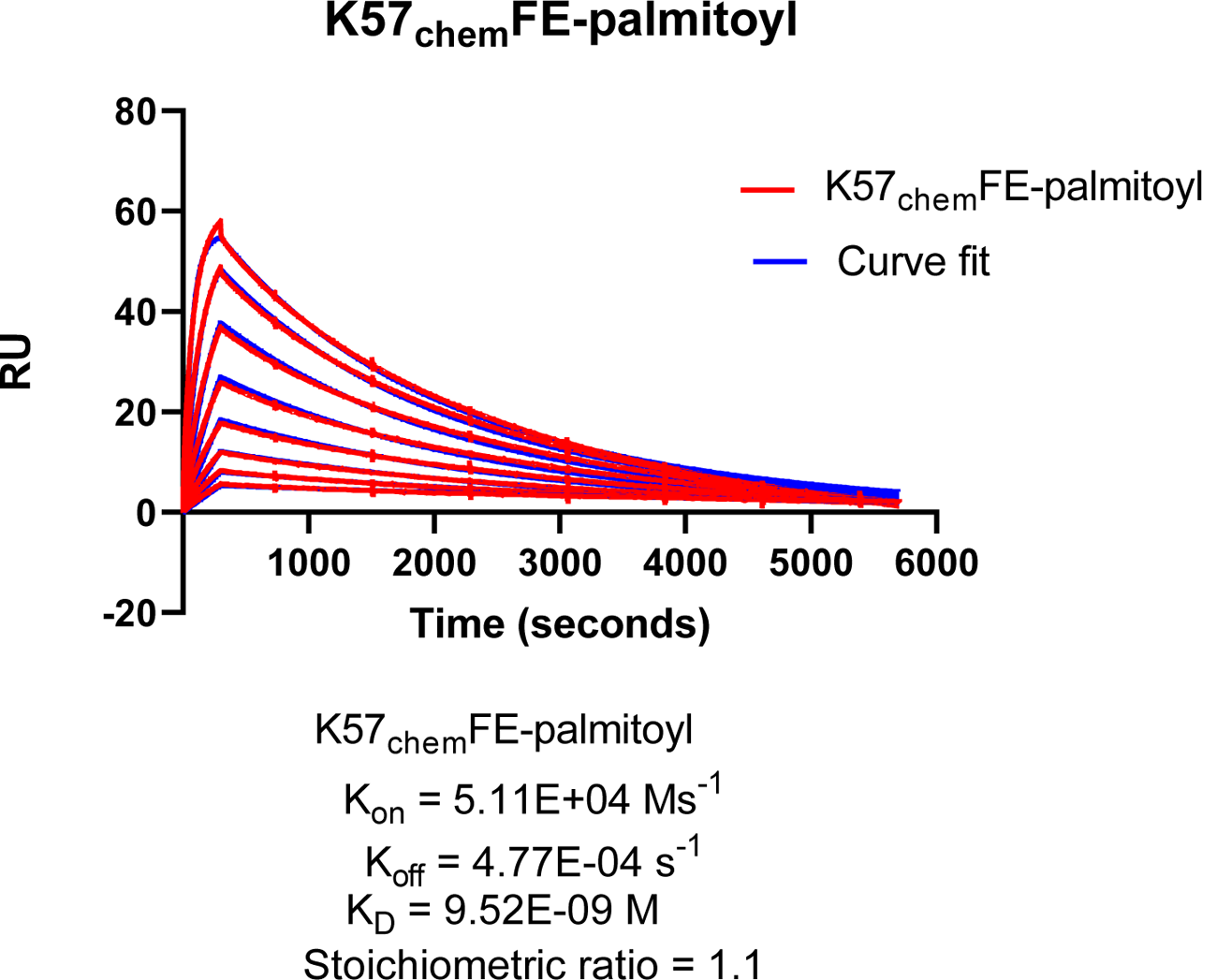
Binding of K57_chem_FE-palmitoyl to human C5 by SPR multi-cycle kinetics.

## References

1. Berens, S. J., Wylie, D. E. & Lopez, O. J. Use of a single VH family and long CDR3s in the variable region of cattle Ig heavy chains. Int Immunol 9, 189–199, doi:10.1093/intimm/9.1.189 (1997).

2. Dong, J., Finn, J. A., Larsen, P. A., Smith, T. P. L. & Crowe, J. E., Jr. Structural Diversity of Ultralong CDRH3s in Seven Bovine Antibody Heavy Chains. Front Immunol 10, 558, doi:10.3389/fimmu.2019.00558 (2019).

3. Stanfield, R. L., Wilson, I. A. & Smider, V. V. Conservation and diversity in the ultralong third heavy-chain complementarity-determining region of bovine antibodies. Sci Immunol 1, doi:10.1126/sciimmunol.aaf7962 (2016).

4. Wang, F. et al. Reshaping antibody diversity. Cell 153, 1379–1393, doi:10.1016/j.cell.2013.04.049 (2013).

5. Stanfield, R. L. et al. The Unusual Genetics and Biochemistry of Bovine Immunoglobulins. Adv Immunol 137, 135–164, doi:10.1016/bs.ai.2017.12.004 (2018).

6. Macpherson, A. et al. Isolation of antigen-specific, disulphide-rich knob domain peptides from bovine antibodies. PLoS Biol 18, e3000821, doi:10.1371/journal.pbio.3000821 (2020).

7. Macpherson, A. et al. The allosteric modulation of complement C5 by knob domain peptides. Elife 10, doi:10.7554/eLife.63586 (2021).

8. Greenberg, A. S. et al. A new antigen receptor gene family that undergoes rearrangement and extensive somatic diversification in sharks. Nature 374, 168–173, doi:10.1038/374168a0 (1995).

9. Hamers-Casterman, C. et al. Naturally occurring antibodies devoid of light chains. Nature 363, 446–448, doi:10.1038/363446a0 (1993).

10. Muyldermans, S., Atarhouch, T., Saldanha, J., Barbosa, J. A. & Hamers, R. Sequence and structure of VH domain from naturally occurring camel heavy chain immunoglobulins lacking light chains. Protein Eng 7, 1129–1135, doi:10.1093/protein/7.9.1129 (1994).

11. Hartmann, L. et al. VHH characterization. Comparison of recombinant with chemically synthesized anti-HER2 VHH. Protein Sci 28, 1865–1879, doi:10.1002/pro.3712 (2019).

12. Martin, W. L., West, A. P., Jr., Gan, L. & Bjorkman, P. J. Crystal structure at 2.8 A of an FcRn/heterodimeric Fc complex: mechanism of pH-dependent binding. Mol Cell 7, 867–877, doi:10.1016/s1097-2765(01)00230-1 (2001).

13. Datta-Mannan, A. Mechanisms Influencing the Pharmacokinetics and Disposition of Monoclonal Antibodies and Peptides. Drug Metab Dispos 47, 1100–1110, doi:10.1124/dmd.119.086488 (2019).

14. Hamley, I. W. PEG-peptide conjugates. Biomacromolecules 15, 1543–1559, doi:10.1021/bm500246w (2014).

15. Wang, J., Shen, D. & Shen, W. C. Preparation, purification, and characterization of a reversibly lipidized desmopressin with potentiated anti-diuretic activity. Pharm Res 16, 1674–1679, doi:10.1023/a:1018929312715 (1999).

16. Craik, D. J. Chemistry. Seamless proteins tie up their loose ends. Science 311, 1563–1564, doi:10.1126/science.1125248 (2006).

17. Gongora-Benitez, M., Tulla-Puche, J. & Albericio, F. Multifaceted roles of disulfide bonds. Peptides as therapeutics. Chem Rev 114, 901–926, doi:10.1021/cr400031z (2014).

18. DePalma, A. Peptides: New processes, lower costs. Genetic Engineering & Biotechnology News 35, 24–26 (2015).

19. Schubert, J. et al. Eculizumab, a terminal complement inhibitor, improves anaemia in patients with paroxysmal nocturnal haemoglobinuria. Br J Haematol 142, 263–272, doi:10.1111/j.1365-2141.2008.07183.x (2008).

20. Roth, A. et al. Ravulizumab (ALXN1210) in patients with paroxysmal nocturnal hemoglobinuria: results of 2 phase 1b/2 studies. Blood Adv 2, 2176–2185, doi:10.1182/bloodadvances.2018020644 (2018).

21. Sampei, Z. et al. Antibody engineering to generate SKY59, a long-acting anti-C5 recycling antibody. PLoS One 13, e0209509, doi:10.1371/journal.pone.0209509 (2018).

22. Ricardo, A. et al. Development of RA101348, a Potent Cyclic Peptide Inhibitor of C5 for Complement-Mediated Diseases. Blood 124, 2936–2936, doi:10.1182/blood.V124.21.2936.2936 (2014).

23. Biesecker, G., Dihel, L., Enney, K. & Bendele, R. A. Derivation of RNA aptamer inhibitors of human complement C5. Immunopharmacology 42, 219–230, doi:10.1016/s0162-3109(99)00020-x (1999).

24. Jendza, K. et al. A small-molecule inhibitor of C5 complement protein. Nature chemical biology 15, 666–668 (2019).

25. Mastellos, D. C., Ricklin, D. & Lambris, J. D. Clinical promise of next-generation complement therapeutics. Nat Rev Drug Discov 18, 707–729, doi:10.1038/s41573-019-0031-6 (2019).

26. Hill, C. P., Yee, J., Selsted, M. E. & Eisenberg, D. Crystal structure of defensin HNP-3, an amphiphilic dimer: mechanisms of membrane permeabilization. Science 251, 1481–1485, doi:10.1126/science.2006422 (1991).

27. Pineda, S. S. et al. Structural venomics reveals evolution of a complex venom by duplication and diversification of an ancient peptide-encoding gene. Proceedings of the National Academy of Sciences 117, 11399–11408, doi:10.1073/pnas.1914536117 (2020).

28. Blanco, M. J. Building upon Nature’s Framework: Overview of Key Strategies Toward Increasing Drug-Like Properties of Natural Product Cyclopeptides and Macrocycles. Methods Mol Biol 2001, 203–233, doi:10.1007/978-1-4939-9504-2_10 (2019).

29. Nicke, A., Ulens, C., Rolland, J.-F. & Tsetlin, V. I. Editorial: From Peptide and Protein Toxins to Ion Channel Structure/Function and Drug Design. Frontiers in Pharmacology 11, doi:10.3389/fphar.2020.548366 (2020).

30. Keshari, R. S. et al. Inhibition of complement C5 protects against organ failure and reduces mortality in a baboon model of Escherichia coli sepsis. Proc Natl Acad Sci U S A 114, E6390–E6399, doi:10.1073/pnas.1706818114 (2017).

31. Ricardo, A. et al. Preclinical Evaluation of RA101495, a Potent Cyclic Peptide Inhibitor of C5 for the Treatment of Paroxysmal Nocturnal Hemoglobinuria. Blood 126, 939–939, doi:10.1182/blood.V126.23.939.939 (2015).

32. Thomas, A. M. et al. Combined Inhibition of C5 and CD14 Attenuates Systemic Inflammation in a Piglet Model of Meconium Aspiration Syndrome. Neonatology 113, 322–330, doi:10.1159/000486542 (2018).

33. Strömberg, P. et al. Development of Affibody® C5 inhibitors for versatile and efficient therapeutic targeting of the terminal complement pathway: 114. Molecular Immunology 61 (2014).

34. Safety and Tolerability of SOBI002 in Healthy Volunteers Following Single and Repeated Administration, <https://ClinicalTrials.gov/show/NCT02083666> (

35. Krissinel, E. & Henrick, K. Inference of macromolecular assemblies from crystalline state. J Mol Biol 372, 774–797, doi:10.1016/j.jmb.2007.05.022 (2007).

36. Chemical Computing Group ULC, S. S. W., Suite #910, Montreal, QC, Canada, H3A 2R7. Molecular Operating Environment (MOE). (2019).

37. Ding, Y. et al. Impact of non-proteinogenic amino acids in the discovery and development of peptide therapeutics. Amino Acids 52, 1207–1226, doi:10.1007/s00726-020-02890-9 (2020).

38. Zhang, R. & Monsma, F. Binding kinetics and mechanism of action: toward the discovery and development of better and best in class drugs. Expert Opin Drug Discov 5, 1023–1029, doi:10.1517/17460441.2010.520700 (2010).

39. Copeland, R. A. The drug–target residence time model: a 10-year retrospective. Nature Reviews Drug Discovery 15, 87–95, doi:10.1038/nrd.2015.18 (2016).

40. Lawson, A. D. G. et al. Modulating Target Protein Biology Through the Re-mapping of Conformational Distributions Using Small Molecules. Frontiers in Chemistry 9, doi:10.3389/fchem.2021.668186 (2021).

41. Lawson, A. D. G., MacCoss, M. & Heer, J. P. Importance of Rigidity in Designing Small Molecule Drugs To Tackle Protein-Protein Interactions (PPIs) through Stabilization of Desired Conformers. J Med Chem 61, 4283–4289, doi:10.1021/acs.jmedchem.7b01120 (2018).

42. Wang, J., Chow, D., Heiati, H. & Shen, W. C. Reversible lipidization for the oral delivery of salmon calcitonin. J Control Release 88, 369–380, doi:10.1016/s0168-3659(03)00008-7 (2003).

43. Hillmen, P. et al. Pegcetacoplan versus Eculizumab in Paroxysmal Nocturnal Hemoglobinuria. N Engl J Med 384, 1028–1037, doi:10.1056/NEJMoa2029073 (2021).

44. Katragadda, M., Magotti, P., Sfyroera, G. & Lambris, J. D. Hydrophobic effect and hydrogen bonds account for the improved activity of a complement inhibitor, compstatin. J Med Chem 49, 4616–4622, doi:10.1021/jm0603419 (2006).

45. Kapoerchan, V. V. et al. Design of azidoproline containing gluten peptides to suppress CD4+ T-cell responses associated with celiac disease. Bioorg Med Chem 16, 2053–2062, doi:10.1016/j.bmc.2007.10.091 (2008).

46. Gentilucci, L., De Marco, R. & Cerisoli, L. Chemical modifications designed to improve peptide stability: incorporation of non-natural amino acids, pseudo-peptide bonds, and cyclization. Curr Pharm Des 16, 3185–3203, doi:10.2174/138161210793292555 (2010).

47. Hallam, T. J., Wold, E., Wahl, A. & Smider, V. V. Antibody conjugates with unnatural amino acids. Mol Pharm 12, 1848–1862, doi:10.1021/acs.molpharmaceut.5b00082 (2015).

48. Brofelth, M. et al. Site-specific photocoupling of pBpa mutated scFv antibodies for use in affinity proteomics. Biochim Biophys Acta Proteins Proteom 1865, 985–996, doi:10.1016/j.bbapap.2017.03.007 (2017).

49. Wang, L., Brock, A., Herberich, B. & Schultz, P. G. Expanding the genetic code of Escherichia coli. Science 292, 498–500, doi:10.1126/science.1060077 (2001).

50. Lindstedt, P. R., Francesco A. Aprile, Pietro Sormanni, Robertinah Rakoto, Christopher M. Dobson, Gonçalo JL Bernardes, and Michele Vendruscolo. Systematic Activity Maturation of a Single-Domain Antibody with Non-canonical Amino Acids through Chemical Mutagenesis. Cell Chemical Biology (2020).

51. Hartrampf, N. et al. Synthesis of proteins by automated flow chemistry. Science 368, 980–987, doi:10.1126/science.abb2491 (2020).

52. Expanding access to monoclonal antibody-based products: A global call to action, <https://www.iavi.org/news-resources/expanding-access-to-monoclonal-antibody-based-products-a-global-call-to-action> (

53. Macpherson, A. et al. The rational design of affinity-attenuated OmCI for the purification of complement C5. J Biol Chem 293, 14112–14121, doi:10.1074/jbc.RA118.004043 (2018).

54. Atherton, E., Atherton, E. & Sheppard, R. C. Solid Phase Peptide Synthesis: A Practical Approach. (IRL Press, 1989).

55. Schrodinger, LLC. The PyMOL Molecular Graphics System, Version 1.8 (2015).

